# Integrated Manufacturing Platform for Cortical Organoids Reveals Early Phosphatase and Prefoldin Dysregulation in *MAPT* V337M Neurons

**DOI:** 10.1101/2023.07.11.548571

**Authors:** Taylor Bertucci, Kathryn R. Bowles, Steven Lotz, Shiraz Bheda, Le Qi, Farhad Farjood, Katherine Whitton, Susan K. Goderie, Susan Borden, Amelia Rossi, Kate Tubbesing, Chiara Pedicone, Laura Maria Oja, Keith Lane, Ryan Lotz, Hailey Lotz, Jack Huber, Rebecca Chowdhury, Shona Joy, Michael Miller, David C. Butler, Brigitte L. Arduini, Heide F B Murray, Carl Alexander Sandhof, M. Catarina Silva, Stephen J. Haggarty, Leander Dony, Fabian J. Theis, Barbara Treutlein, Celeste M. Karch, Daniel H. Geschwind, Alison M. Goate, Jeffrey Stern, Sally Temple

## Abstract

Human pluripotent stem cell (hPSC)-derived cortical organoids are powerful models but are often limited by low efficiency, variability, and stress-related artifacts. To address these challenges, we developed a scalable organoid platform with end-to-end quality control (QC) metrics spanning manufacturing and single-cell RNA-sequencing (scRNA-seq), developed using eight *MAPT* mutation isogenic line sets relevant to frontotemporal dementia (FTD-tau). Using a 96 slit-well format, we achieved ∼100% production efficiency across 64 lines. Controlled-release FGF2 enhanced iPSC pluripotency and reduced mesendodermal contaminants, while optimized SB431542 dosing enhanced cortical patterning across lines with variable *TGFBR1/ALK5* expression. The resulting organoids displayed transcriptomic profiles and low-stress signatures closely aligned with the developing human cortex. Applying a cortical organoid scRNA-seq index (CortiCOSI), we identified early dysregulation of phosphatase regulators (*PPP2CA, ANP32A*) and the prefoldin subunit *PFDN6* in *MAPT* V337M excitatory neurons before tau hyperphosphorylation and oligomerization. This platform improves scalability, reproducibility, and mechanistic insight in cortical organoid studies.

## Introduction

hPSC-derived brain organoids patterned toward the cerebral cortex are valuable models that capture the 3D multicellular interactions occurring *in vivo* in cortical tissue.^1–10^ During cortical development, neural progenitor cells, including ventricular radial glia, outer radial glia, and intermediate progenitor cells, generate first deep-layer and then upper-layer cortical excitatory neurons, inhibitory neurons, astrocytes, and oligodendrocytes. hPSC-derived cortical organoids recapitulate this developmental sequence of *in vivo* cell production.^4,11–17^ The resulting organoids can be cultured long-term, enabling temporal studies of cortical development and disease progression.^16^ ^18–21^

hPSC-derived cortical organoids can model various neurological and psychiatric conditions, including neurodevelopmental and neurodegenerative diseases.^20,22–30^ For example, we and others have used cortical organoids to study the pathological processes underlying dominantly inherited frontotemporal dementia (FTD) caused by mutations in *MAPT,* the gene encoding tau. FTD encompasses a clinical spectrum of behavioral, cognitive, language, and movement disorders. In individuals with *MAPT* mutations, the underlying neuropathology is characterized by frontotemporal lobular degeneration with tau pathology (FTLD-tau).^18,19,30–34^ Over 50 disease-causing mutations in the *MAPT* gene have been identified.^35,36^ ^37^

We previously characterized the time course of changes due to the *MAPT* V337M mutation.^18^ Early changes detected in 2-month organoids include altered expression of the RNA-binding protein *ELAVL4*, disrupted alternative splicing, premature development, disruption of glutamatergic pathways, and altered proteostasis.^18^ Later-stage organoids exhibited accumulated abnormally phosphorylated tau, and selective glutamatergic cortical neuron death, which occurred predominantly at six months.^18^ Thus, cortical organoids can recapitulate key temporal features of FTLD-tau pathogenesis and enable identification of novel genes and pathways. This prompted us to expand our analysis to additional *MAPT* mutation lines from the Tau Consortium collection^38^ to identify FTLD-associated pathophysiological phenotypes. However, cortical organoid modeling across large line sets from multiple donors is hindered by high organoid variability and elevated stress signatures, which limit reproducibility and effective disease modeling.^4,19,39–44^ These challenges likely reflect intrinsic differences between hPSC lines, both between donors and within donors,^42,45,46^ such that applying uniform protocols generates heterogeneous organoid outcomes.^13,42^ Thos variability introduces noise and obscures genotype-driven phenotypes, creating a critical need for protocols that reproducibly generate well-patterned, low-stress cortical organoids across lines with quantifiable QC metrics. This need is particularly acute for collections derived from patients with rare mutations, such as those driving FTLD-tau, where opportunities to generate additional lines is limited.

Here, we report a scalable cortical organoid protocol with defined end-to-end QC metrics spanning manufacturing and scRNA-seq, developed using 17 lines, including 7 *MAPT* mutation donors. Well-patterned organoids generated using this platform exhibited cortical-specific transcriptomic signatures, consistent patterning, and markedly reduced cellular stress signatures. These features enabled the development of CortiCOSI, a composite metric for defining well-patterned cortical organoids in scRNA-seq samples, also applicable to datasets generated by alternative methods. We applied CortiCOSI to our prior V337M scRNA-seq dataset, for which extensive phenotypic data are available, and identified isogenic pairs with consistent, well-patterned profiles. Transcriptomic analysis of this CortiCOSI-selected subset revealed early molecular signatures undetectable in the full dataset, implicating reduced PP2A-mediated dephosphorylation and diminished prefoldin complex function at 2 months as drivers of tau hyperphosphorylation and oligomeric tau accumulation at later time points. These findings demonstrate the power of consistent, QC-driven human organoid modeling to identify early transcriptomic changes and establish a tractable system for mechanistic studies and future analyses across multiple *MAPT* mutation lines.

## Results

### Improved efficiency and consistency using a 96 slit-well plate format

We sought to generate large-scale batches of hPSC-derived cortical organoids to enable phenotypic comparisons between *MAPT* mutation lines versus isogenic controls. We initially used a feeder-free hPSC organoid protocol,^6^ summarized in **Figure 1A**. In this dish-based method, organoids frequently fused, necessitating manual cutting to separate them. This is laborious, introduces variable levels of damage, and produces organoids of different sizes, particularly at early stages when regional specification is occurring. Even with manual cutting, the protocol is only approximately 10% efficient. Thus, to improve scalability, efficiency, and consistency, we translated this protocol into a 96 slit-well plate format with shared medium between the wells and applied the identical media sequence as described in the dish method (**Figure 1A,B**). This configuration physically isolates individual organoids to prevent organoid fusion while maintaining uniform culture conditions.

**Figure 1.**
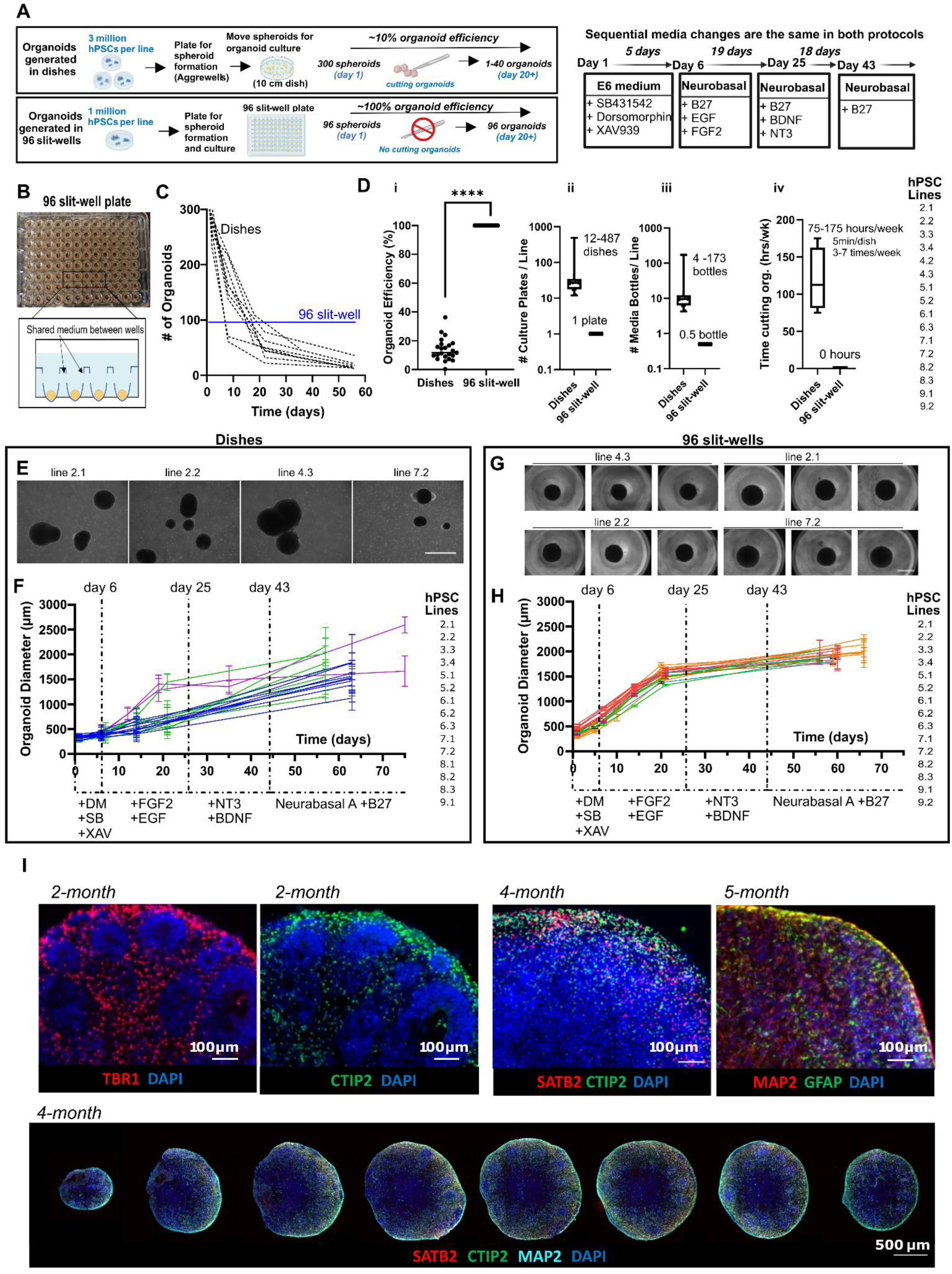
Efficient and consistent organoid generation in a 96 slit-well plate format. **A, B)** Schematics comparing the original dish-based protocol with the 96 slit-well plate (slit-well protocol_v1); medium is shared across slit-wells. **C)** Organoid counts over time in dishes (dotted lines) versus 96 slit-well plates (purple line); 7 lines (2 repeated, giving 9 samples/condition) pooled from 2 independent experiments. **D)** Efficiency: (i) Number of organoids remaining at day 20 divided by the number of day 1 hPSC spheroids for dish versus 96 slit-well methods; 16 lines (7 repeated twice; 23 samples/condition) pooled from 3 independent experiments; unpaired two-tailed t-test, ****p<0.00005. (ii–iv) Comparison of plasticware, culture medium consumption, and hands-on time required to generate 96 two-month-old organoids in dishes versus 96 slit-well plates; calculated based on the manufacturing efficiency of 16 lines from 8 donors (**see 1D**). **E, F)** Dish-based method: (E) representative phase images of 14-day organoids showing heterogeneity (scale bar, 200 µm). (F) Growth curves (mean diameter/line) show variable growth; error bars, standard deviation (SD); 15 lines (4 lines repeated twice; 19 total samples), 4–12 organoids/line/time point, pooled from 3 independent experiments (indicated by color). **G, H)** Slit-well method: (G) representative phase images of 20-day organoids showing consistency (scale bar, 200 µm). (H) Growth curves (mean diameter/line) are uniform; error bars, SD; 16 lines (6 repeated 2–3 times; 32 total samples), 4–12 organoids/line/time point, pooled from 4 independent experiments (indicated by color). **I)** Upper image panels: Well-patterned cortical organoids were generated and maintained in 96 slit-well plates for 2-4 months, then immunostained for cortical markers. Deep-layer TBR1+ and CTIP2+ cortical neurons emerge by 2 months; upper-layer SATB2+ neurons and GFAP+ astrocytes are robust by 4-5 months; lines 3.4, 5.1. Lower image panel: Serial sections from a single organoid demonstrate cortical patterning throughout; line 3.3. See also Supplemental **Figure S1** and **Table S1**.

hPSC spheroids formed in slit-wells showed significantly higher expression of the pluripotency-associated genes *POU5F1/OCT4, NANOG,* and *SOX2* compared to spheroids formed in Aggrewells, and more consistent expression of *PAX6,* a marker of early neural and forebrain specification, at day 6 (**Figure S1A**). Organoid production efficiency approached 100%, with reduced reagent usage and labor, eliminating the need for manual cutting (**Figure 1C, D**). Compared with dishes, individual organoids generated in 96-well slit plates were more consistent in size, shape, growth pattern, and gene expression across lines and experiments (**Figure 1E-H; Figure S1B-D).** Growth in conventional 96-well plates, which typically requires transfer to a different culture format for longer-term culture,^47,48^ did not produce as consistent an early organoid profile as observed in the slit-well plates (**Figure S1Aii**).

While the 96 slit-well format improved efficiency and growth consistency, it did not resolve variability in cortical patterning across lines. We generated organoids from 8 isogenic sets (n=16 lines) (**Table S1**), with each line cultured in a separate 96 slit-well plate. Only 6 out of the 16 lines (37%) robustly formed PAX6+FOXG1+ rosettes of forebrain progenitors and produced neurons expressing the deep-layer cortical markers CTIP2/BCL11B and TBR1 at 2-months (**Figure S1E,F**). In successful cases, the well-patterned organoids continued to mature in slit-well plates and, by 4 months, included neurons expressing the upper-layer marker SATB2 together with GFAP+ astrocytes (**Figure 1I**). The variability in patterning success across lines motivated further protocol optimization. We therefore investigated factors influencing the earliest stages of differentiation, beginning with the pluripotency status of the starting hPSC lines.

### Robust hPSC pluripotency is critical for successful cerebral cortical organoid patterning

FGF2, a key regulator maintaining hPSC pluripotency, is thermally unstable, and even daily feeding with established media such as mTeSR1 results in periods of low FGF2 concentration.^49–51^ Insufficient FGF2 signaling can compromise pluripotency, promote spontaneous differentiation and increase off-target lineage specification. Because thermostable or hyperstable versions differ biologically from native FGF2,^52,53^ we instead evaluated a sustained-release formulation of native FGF2 (FGF2DISC). We compared this approach with standard daily feeding using mTeSR1, which contains soluble native FGF2^54^ (**Figure 2A**).

**Figure 2.**
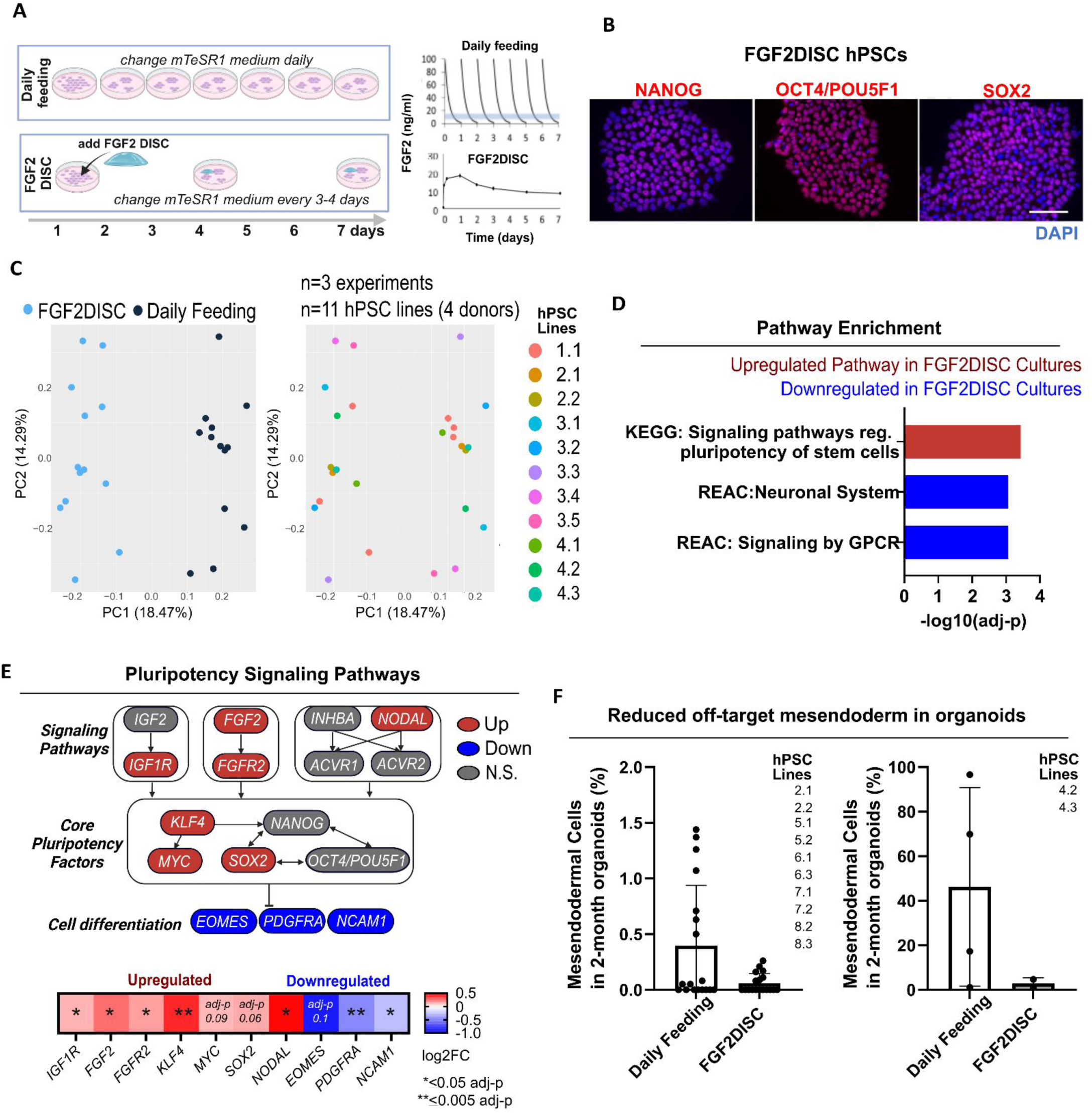
Sustained-release FGF2 increases pluripotency of hPSCs. **A**) i) Schematic showing daily mTeSR1 feeding versus twice-a-week mTeSR1 feeding plus FGF2DISCs. ii) Top panel: schematic showing the fluctuating FGF2 levels when medium containing 100 ng/ml soluble native FGF2 (e.g., mTeSR1) is added daily, based on a measured FGF2 half-life of 4 hours;^22^ Blue line indicates the ideal FGF2 concentration. Lower panel: FGF2 release over time from an FGF2DISC at 37°C assessed by flow cytometry, n = 3 DISC samples, error bars indicate SD. **B**) hPSC were cultured with FGF2DISCs and immunostained for pluripotency markers; line 1.1; scale bar=100 µm. **C-E**) Bulk RNA-seq comparing FGF2DISC cultures to daily feeding; 11 lines from 4 donors, one line repeated in each experiment as a standard (13 samples per condition), data compiled from 3 independent experiments. **C**) PCA plots colored by condition (left), hPSC line (right); key of sample codes. **D**) The top-ranked significantly enriched pathways in FGF2DISC cultures identified with g:Profiler2; REAC=Reactome. **E)** Diagram and heatmap of gene changes in pluripotency signaling networks in FGF2DISC cultures; N.S=not significant. **F**) Bar graphs showing the percentage of mesendodermal cells in 2-month organoids: i) 18 samples per condition (9 lines from 2 independent experiments); ii) Donor #4 organoids shown separately (2 lines). See also Supplemental **Figure S2, S3** and **Table S2-S5.**

Eleven hPSC lines from four donors were exposed to the two different feeding regimens for 3-4 weeks and then assessed by bulk RNA-seq. iPSCs grown with FGF2DISCs showed robust expression of pluripotency markers (**Figure 2B**). PCA analysis revealed that the feeding regimen was the primary driver of transcriptomic variability (**Figure 2C**). Differential expression analysis and subsequent pathway enrichment analysis identified the most enriched pathway associated with FGF2DISCs as “signaling pathways regulating pluripotency of stem cells” (**Figure 2D**, **Tables S2, S3**). Nodal/TGFβ, FGF2 signaling factors, and transcription factors with critical roles in maintaining pluripotency^5,51^ were upregulated with FGF2DISC culture, while cell differentiation pathways and related genes were significantly downregulated, including *EOMES, PDGFRA,* and *NCAM1* (**Figure 2E**). Gene expression variability between samples (across lines and experimental batches), quantified from raw log counts per million (CPM) and evaluated using standard deviation (SD), Euclidean distance, or Spearman’s rho, was reduced with FGF2DISCs (**Figure S2A**), suggesting that some lines are particularly susceptible to fluctuating levels of FGF2. Indeed, an isogenic set of lines from hPSC donor #4 (three lines), which showed the lowest pluripotency profiles in daily mTeSR1 culture, expressed significantly lower endogenous *FGF2* than other donor lines at the pluripotent stage, perhaps accounting for their susceptibility (**Figure S2B, C**). These lines, when grown with daily mTeSR1 feeding, produced organoids with abnormal growth curves, low neuronal content, and increased proportions of mesendodermal cell types, characterized by expression of genes such as *COL1A2, TMEM119*, and *DCN* (**Figure S3A-D, Table S4**).

While donor #4 organoids provided the most robust example, off-target mesendodermal cell types were detectable across donors at lower levels. In organoids derived from daily-mTeSR1 fed hPSC cultures, mesendodermal cells comprised 0.5-1.5% in approximately one-third of the samples, excluding donor #4 (7/18), and reached up to 96% in donor #4 organoids (**Figure 2F**). In contrast, organoids generated from FGF2DISC hPSC cultures showed a marked reduction in these off-target populations: excluding donor #4, no sample (0/18) had levels above 0.5%, while the incidence in donor #4 was reduced to < 5% (**Figure 2F**). Thus, optimizing hPSC culture conditions improved neural induction and reduced mesendodermal contamination. However, enhancing pluripotency alone did not fully account for variability in cortical fate acquisition across donors, prompting an investigation into early patterning cues.

### Tailoring SB431542 concentration improved cortical patterning in hPSC lines with different TGFBR1/ALK5 expression

To pattern the cerebral cortex, three primary pathways are inhibited during the first 6 days of differentiation using a small-molecule cocktail: dorsomorphin (DM) to block BMP type I receptors (ALK2/3/6), SB431542 to inhibit Lefty/Activin/TGF-beta signaling via ALK4/5/7, and XAV939 to suppress the WNT pathway. We hypothesized that donor-specific variability could be due to differing baseline requirements for these cortical patterning molecules. Indeed, hPSC lines that failed in the slit-well protocol_v1 (i.e., daily feeding hPSC cultures and patterning factors established in the dish method^6^, **Figure 1A**) exhibited altered baseline expression of genes associated with the TGF-beta, BMP, and WNT signaling pathways (**Table S5**). Hence, we manipulated the levels of these patterning factors and analyzed the outcomes.

The slit-well protocol_v1 (**Figure 1A**) and the dish-based protocol^6^ both used 2.5 µM DM. Testing higher concentrations of DM or another BMP inhibitor, LDN193189, did not improve *FOXG1* expression at day 20, and organoids lacked TBR1+ and CTIP2/BCL11B+ cells at two months, despite robust MAP2 neuronal staining (**Figure S4A**). Increasing the XAV939 concentration to 5 µM or substituting 2 µM IWR, an alternative WNT pathway inhibitor, generally compromised organoid health, with only modest benefits in cortical progenitor and neuronal gene expression in a subset of lines (**Figure S4B, C**). In contrast, varying SB431542 concentrations from the original 10 µM up to 30 µM significantly impacted expression of the cortical progenitor markers *PAX6* and *FOXG1* at 20 days, and optimizing these levels markedly improved cortical patterning (**Figure 3A, S4C-F**). Notably, several hPSC lines that require higher concentrations of SB431542 for optimal cortical patterning exhibited elevated baseline expression of its primary target, *ALK5*, at the pluripotent stage (**Figure S4G**).

**Figure 3.**
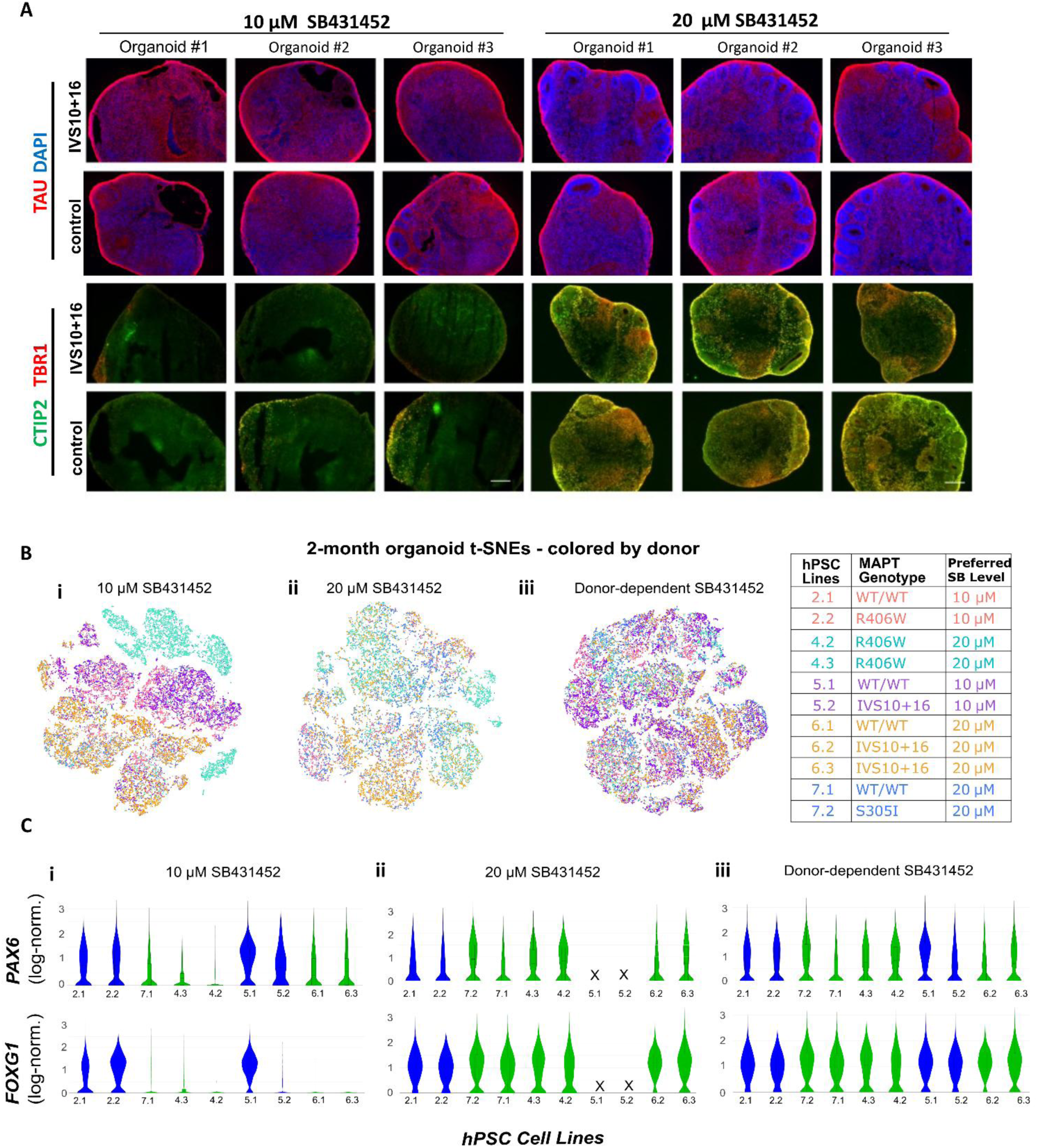
Optimizing the SB431542 concentration (d1-d6) improves the consistency of cortical patterning. **A)** Representative fluorescence images of 2-month-old organoid sections from donor #6 that did not pattern well at 10 μM SB431542 showing they do so when patterned with 20 μM SB431542; organoids in both conditions show robust neuron production, indicated by tau immunostaining (top panel), but have improved expression of cortical deep-layer markers CTIP2 and TBR1 (bottom panel) with 20 µM SB431542; scale bar = 200 µm. **B, C**) scRNA-seq data from 2-month-old organoids (3 organoids pooled/sample) show most consistency when the SB431542 is tailored to each line (10 µM or 20 µM); <u>10 µM treatment:</u> 1-2 lines/donor, 5 donors, (9 samples); <u>20 µM treatment</u>: 2 lines/donor, 5 donors (10 samples). <u>Donor-dependent</u>: 2 lines/donor, 5 donors, 2 lines repeated (12 samples). **B**) t-SNE plots colored by donor and line codes. **D)** Violin plots of *PAX6* and *FOXG1* expression by hPSC line colored based on optimal SB431542 level, blue = 10 µM, green = 20 µM; ‘X’ = organoids did not survive. See also Supplemental **Figure S4.**

scRNA-seq was used to evaluate organoids generated from isogenic pairs of lines from five donors, exposed to either 10 or 20 µM SB431542 during the patterning phase of the protocol (days 1- 6) (**Figure 3B,C**). When organoids were produced using the donor-optimized SB431542 concentration, transcriptomic profiles became more consistent, t-SNE plots showed greater overlap (**Figure 3Bi-iii, 3Ci-iii**), and the coefficient of variation for *FOXG1* expression improved 3.4-fold, decreasing from 107.3% to 32%. Thus, tailored SB431542 dosing improves the consistency of cortical organoid patterning across diverse hPSC donors.

Together, the incorporation of FGF2DISC culture conditions and the optimization of early patterning through tailored SB431542 dosing defined our second-generation slit-well protocol (slit-well protocol_v2) and markedly improved the success rate of well-patterned organoids across hPSC lines. However, achieving high reproducibility requires not only optimized protocol parameters, but also a systematic framework for monitoring organoid quality throughout manufacturing.

### End-to-End QC metrics for cortical organoid manufacture

Our slit-well protocol_v2 and the manufacturing QC platform we have developed, summarized in **Figure 4A,B**, were generated by correlating data collected at different stages of manufacturing to the production of well-patterned 2-month organoids that robustly expressed cortical neuron markers, assessed by IHC. Comprehensive QC for successful cortical organoid production begins at the hPSC stage when 90-100% of cells should be double-positive for SSEA4/TRA-1-60 (**Figure 4B,Ci**). The initial diameter of the slit-well spheroid on day 1, optimally 300-500 μm, is important for proper organoid growth and cortical patterning (**Figure 4B,D**). At day 20, we implement a ‘fail-early’ strategy using qPCR as an in-process checkpoint. *COL1A2* serves as a validated negative QC marker, enabling early detection of off-target mesendoderm contamination, which is associated with reduced pluripotency in the starting hPSC cultures (**Figure 4Cii**). Conversely, the expression levels of the forebrain progenitor markers *PAX6* and *FOXG1* at 20 days are positive predictors (**Figure 4Ei**) that correlate with qPCR levels of the cortical neuron markers *TBR1* and *BCL11B/CTIP2* at two months (**Figure 4Eii,iii**). This is confirmed by sectioning the 2-month-old organoids and demonstrating positive IHC staining for TBR1 and BCL11B/CTIP2 neuronal markers. While no single QC metric is definitive, this integrated QC framework, developed through extensive empirical optimization and validation across diverse experimental conditions and hPSC lines, provides high confidence in the consistency of 2-month cortical organoids.

**Figure 4.**
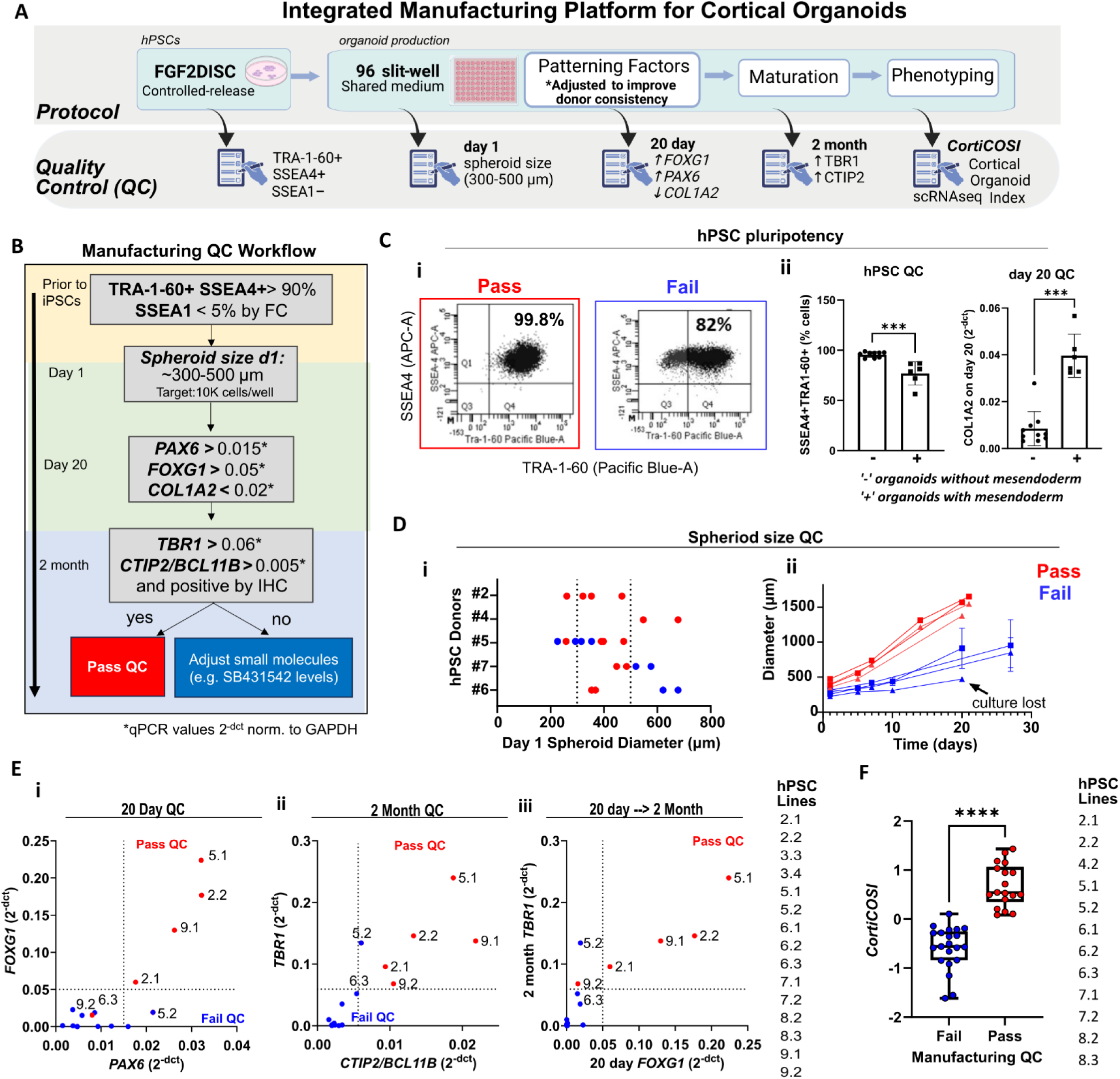
Established end-to-end QC assessments ensure consistent, well-patterned cortical organoids during manufacturing and in scRNA-seq datasets. **A,B)** Schematics of the integrated manufacturing platform for cortical organoids in 96 slit-well plates. **C)** i) hPSC QC metrics: percentage of SSEA4/TRA-1-60 double-positive cells assessed by flow cytometry; see **Figure S2D** for gating strategy and controls. ii) Lower SSEA1+TRA1-60+ in starting hPSCs and higher *COL1A2* expression at 20 day predict off-target mesendodermal cells in 2-month organoids; 12 lines, 16 samples; Mann-Whitney U test (two-tailed), ***p<0.005. **D)** i) Optimal day 1 spheroid size range to achieve successful growth and patterning. ii) Growth curves from organoids from undersized or appropriately sized starting spheroids; data represent means +/- SD of 4–12 organoids; lines 5.1, 5.2, 4 experiments. **E)** Example of day 20 qPCR thresholding strategy. Note *FOXG1* and *PAX6* levels at 20 days by qPCR (i) predict 2-month *TBR1* and *CTIP2* levels (ii, iii). **F**) CortiCOSI for organoids that pass or fail manufacturing QC metrics; 12 lines, 4 experiments; see **Figure S5** for CortiCOSI calculations; unpaired two-tailed t-test, ****p <0.0001. See also **Supplemental Figure S5, S6** and **Table S6.**

Following our work to improve the wet-lab protocol, we expanded the platform to include data-driven validation against developing-brain reference datasets and computational approaches to enhance organoid standardization. We developed an scRNA-seq QC metric to assess cortical organoid quality and consistency. Each sample (using all cells) was evaluated for cortical regional (CR) identity using VoxHunt, benchmarked against both the Allen Brain Atlas (ABA) mouse^39^ and the human BrainSpan^55^ datasets (**Figure S5Ai,ii, Table S6)**. The CR identity value was then integrated with the expression levels of *FOXG1* and *COL1A2*, two genes selected for their relevance to the quality and consistency of cortical organoid manufacturing (**Figure S5Aiii, iv**) to produce a composite z-score (CortiCOSI) that effectively discriminated pass and fail QC groups defined by manufacturing QC with minimal overlap (**Figure 4F, S5B**). Although Z-scores are relative, integrating these three orthogonal benchmarks into a single CortiCOSI metric enabled visualization and comparison of inter-sample variation across the scRNA-seq dataset.

### QC-validated organoids exhibit authentic cortical identity and low stress

To evaluate the fidelity and physiological state of QC-pass manufactured organoids, we assessed CR identities and cellular stress using single-cell transcriptomic analysis. Individual cells were annotated using SingleR, referencing developing human cortex datasets, following our previous methodology.^18^ Cells annotated as glutamatergic excitatory neurons were identified and evaluated for cortical regional identity values using mouse^39^ and human^55^ datasets. Organoids that passed manufacturing QC were enriched for gene expression signatures characteristic of authentic cortical excitatory neurons, whereas failed organoids displayed signatures associated with diverse non-cortical brain regions (**Figure S6A,B**).

Previous analyses of cortical organoids generated by other methods have reported high glycolytic and endothelium reticulum (ER) stress signatures, which may interfere with cell specification and limit disease modeling.^4,42–44^ Compared with datasets from the Human Neural Organoid Cell Atlas (HNOCA)^56^ and primary developing human brain samples,^11,13,57^ well-patterned wild-type (WT) organoids generated using our platform contained neurons with close similarity to primary tissue and the lowest stress marker expression among cortical-guided organoid studies that used more than one line (**Figure 5A, Table S7,S8**). Importantly, this effect persisted after controlling for age: comparison of our 60-day organoids with other datasets spanning 30-90 days showed that reduced stress signatures were not associated with age of organoids or primary tissue (**Figure S7A-C**).

**Figure 5.**
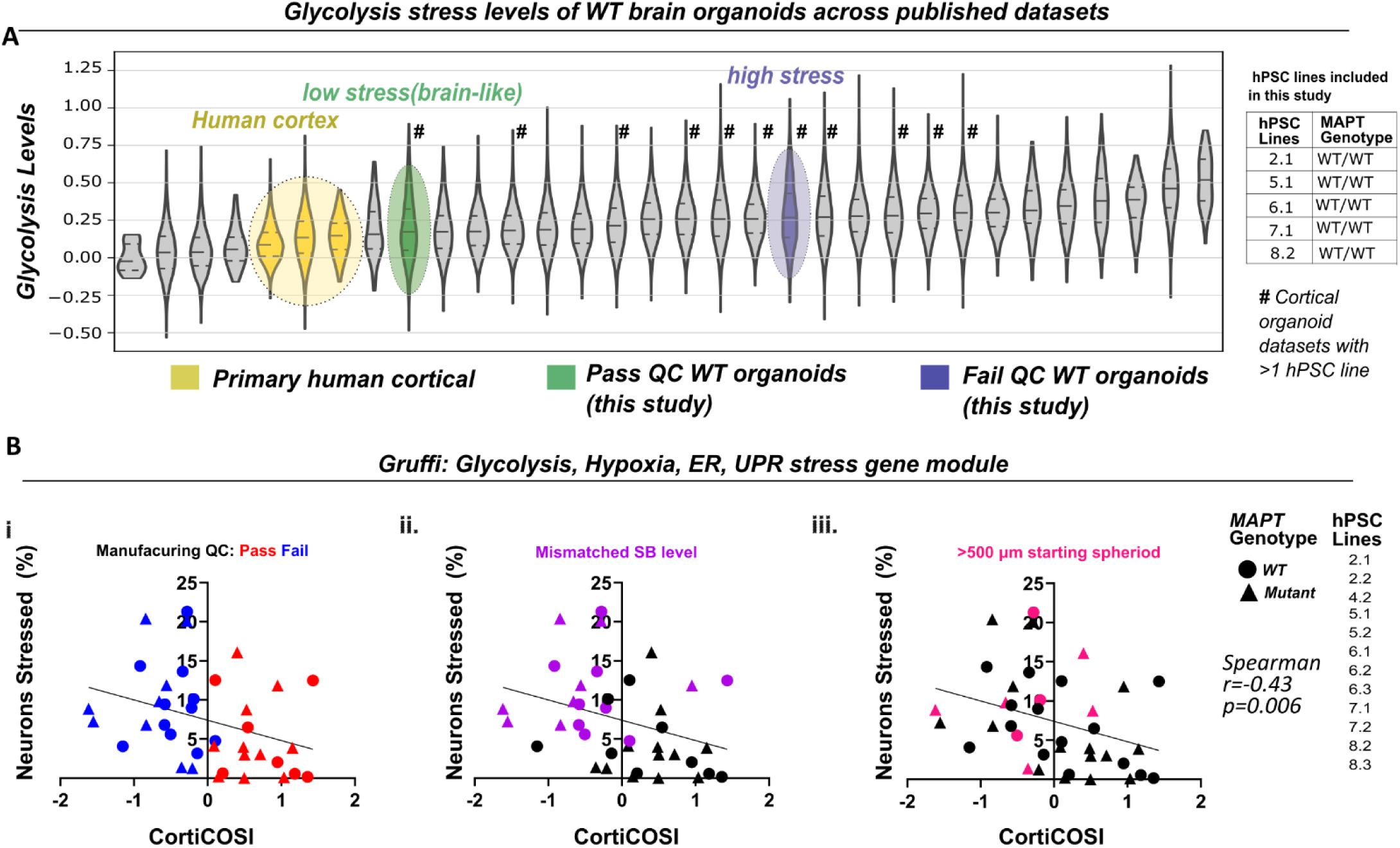
Well-patterned WT cortical organoids identified using QC metrics demonstrate reduced cell stress signatures similar to developing human cortex. **A)** Metabolic neuron stress values across HNOCA organoid datasets^1^, human cortical tissue datasets^2–4^, and datasets generated in this study, stratified by pass/fail status based on QC manufacturing metrics; see **Table S7** for genes included in glycolysis gene score; see **Table S8** for detailed sample information; Lines within violin plots indicate quartiles of the respective distributions. **B)** Percentage of stressed neurons, quantified by Gruffi, plotted against CortiCOSI scores; see **Figure S5** for CortiCOSI calculation. Organoids generated within quality manufacturing metrics, Pass QC, donor-preferred SB431542 levels, and recommended spheroid size exhibit the lowest stress levels and highest patterning fidelity; Spearman’s correlation between neuronal stress and CortiCOSI scores, r= -0.43, p = 0.006. Line indicates linear regression (slope significantly different from 0), p = 0.038. See also Supplemental **Figure S7** and **Tables S7, S8.**

In a complementary assessment of stress, we employed Gruffi, a bioinformatic tool to identify stressed cells in brain organoids based on glycolysis, hypoxia, ER stress, and unfolded protein response (UPR) gene modules.^58^ The proportion of stressed neurons quantified by Gruffi was compared with CortiCOSI scores for each sample (**Figure 5B**). Higher CortiCOSI scores correlated with lower proportions of stressed neurons, whereas organoids with lower CortiCOSI scores, for example when the SB431542 level or starting spheroid size were not controlled, showed variable but often elevated neuronal stress levels (**Figure 5Bi-iii**), averaging approximately 10%, consistent with reported values from other published organoid datasets.^58^ In contrast, WT organoids manufactured with our improved platform that passed QC contained very few stressed neurons (approximately 0–1%) (**Figure 5Bi**). Together, these data demonstrate that end-to-end quality manufacturing generates well-patterned organoids with reduced cellular stress and transcriptomic profiles more closely aligned with normal cortical development.

### CortiCOSI-enabled detection of early phosphatase and prefoldin dysregulation in MAPT V337M organoids

We next asked whether CortiCOSI could identify high-quality, well-patterned organoids produced using different manufacturing approaches. To test this, we evaluated our previously published dataset of *MAPT* V337M 2-month organoids generated in dishes from daily-fed hPSC cultures, which includes three isogenic sets across seven lines and three batches.^18^ 14% (3/21) of samples exhibited elevated *COL1A2* expression (**Figure S8Ai**). *FOXG1* expression varied (0.28-1.8) as did the CR identity values (mouse reference: 0.53-0.64; human reference: 0.44-0.53) (**Figure S8Aii-iv**). Composite CortiCOSI scores revealed variable concordance among isogenic pairs, with some pairs showing high consistency and others substantial variability (**Figure S8B**). Hence, applying CortiCOSI identified a subset of highly consistent, well-patterned isogenic pairs within the dataset. We then performed paired differential gene expression analysis of TBR1+ excitatory neurons comparing V337M mutant and isogenic CRISPR-corrected V337V (WT) organoids using 1) CortiCOSI-selected high-quality pairs only (**Figure 6A-C**, **Table S9**) and 2) all isogenic pairs, irrespective of organoid quality metrics (**Figure 6A-C**, **Table S10**).

**Figure 6.**
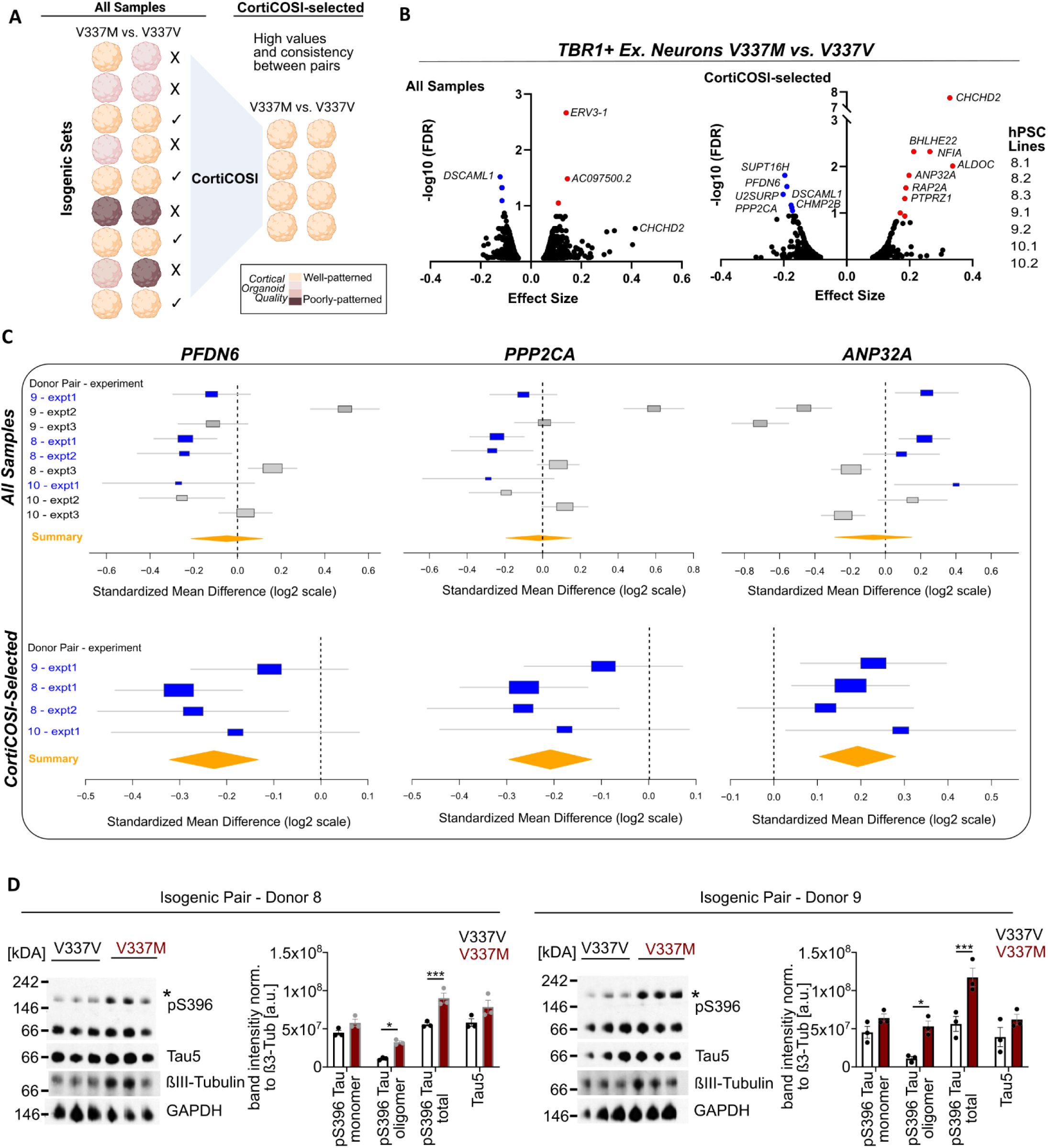
CortiCOSI-selected reanalysis improves detection of differential gene expression in cortical organoids and reveals tau pathology-related perturbations in *MAPT* V337M TBR1+ neurons. **A)** Schematic of the selection method for well-patterned and quality-manufactured samples for CortiCOSI-selected isogenic pair analyses; see also Supplemental **Figure S8**. **B)** Volcano plot of upregulated and downregulated genes in TBR1+ *MAPT* V337M neurons in 2-month organoids vs isogenic controls; i) paired analysis of all samples without assessment of cortical QC metrics or ii) CortiCOSI-selected isogenic pairs. **C)** Forest plots illustrate the consistency of the top differentially expressed genes across CortiCOSI-selected isogenic pairs (bottom) compared to all isogenic pairs (top); Boxes indicate the estimated log2FC for each isogenic pair, lines indicate 95% confidence intervals, vertical dotted line denotes no change. **D)** Western blots and densitometry quantification separated by Blue Native (BN)-PAGE of individual 250-day-old organoids from 4 lines (2 isogenic pairs) of *MAPT* V337M and isogenic V337V control lines, 3 organoids/line, (12 samples of individual organoids); * indicates the multimeric band; bar graphs show mean ± SEM; two-way ANOVA with Holm-Sidaks adjustment for multiple comparisons, *p ≤ 0.05, *** p ≤ 0.001. A.U.=arbitrary units. See also **Figure S9B** for raw blots. See also Supplemental **Figure S8** and **Table S9, S10**

Most notably, TBR1+ neurons showed more differentially expressed genes (DEGs) in the CortiCOSI-selected analysis than in the analysis of all samples (**Figure 6B**). Among the top 50 ranked DEGs, only 2% overlapped between both analyses. These included *CHCHD2*, associated with oxidative phosphorylation and apoptosis (effect size (ES) 0.33; FDR 2.4E-08), and the cell adhesion molecule *DSCAML1* (ES -0.16; FDR 0.12) (**Figure 6B**). Genes uniquely identified in the CortiCOSI-selected comparison included increased expression of the glycolysis-associated gene *ALDOC* (ES 0.34; FDR 0.01), and reduced expression of the splicing regulator *U2SURP* (ES -0.17; FDR 0.076), the chromatin factor *SUPT16H* (ES -0.20; FDR 0.01), and *CHMP2B* (ES -0.17; FDR 0.14), a component of the ESCRT complex. Missense variants of *SUPT16H,* a subunit of the FACT (Facilitates Chromatin Transcription) complex, are linked to neurodevelopmental disorders, including intellectual disabilities, seizures, and corpus callosum abnormalities.^59,60^ Mutations in *CHMP2B* can cause FTD, likely by disrupting ESCRT functions,^61^ and can synergize with the loss of *CHMP2A* to promote tau aggregation.^62^

CortiCOSI-selected pair analysis also identified coordinated dysregulation of PP2A signaling, with decreased expression of the catalytic subunit *PPP2CA* (ES -0.17; FDR 0.015) and increased expression of its endogenous inhibitor *ANP32A* (ES 0.198; FDR 0.1) (**Figure 6B,C**). PP2A signaling is responsible for approximately 70% of total tau phosphatase activity in the human brain,^63^ and its inhibition promotes tau phosphorylation at multiple sites, including Ser-199, Ser-202, Thr-205, Ser-396, and Ser-404.^64^ Consistent with this, we previously reported elevated tau phosphorylation at Ser-202 and Ser-396 V337M organoids at 4 and 6 months,^18^ which aligns with tau pathology observed in V337M patient brain tissue.^65^

The prefoldin complex subunit *PFDN6* (ES -0.2; FDR 0.04) was also among the top-downregulated genes identified in the CortiCOSI-selected analysis (**Figure 6B,C**). The prefoldin complex functions as a molecular chaperone that supports proper folding of cytoskeletal proteins, and impaired prefoldin activity drives early formation of toxic oligomers.^66,67^ Consistent with this possibility, V337M mutant organoids at 8 months exhibited a higher molecular weight oligomeric p-Ser396 tau^68^ band (**Figure 6D**). We therefore speculate that reduced prefoldin complex function may contribute to tau oligomerization. These observations demonstrate that QC-guided organoid stratification enables detection of genotype-associated transcriptional changes that are otherwise obscured by organoid variability.

## Discussion

To define phenotypes associated with *MAPT* mutations in hPSC-derived cortical organoids, we developed a manufacturing platform incorporating enhanced hPSC maintenance, a 96 slit-well culture format, refined patterning conditions, and end-to-end QC metrics. Together, these advances increased the sensitivity for detecting genotype-associated phenotypes. Well-patterned cortical organoids defined by manufacturing QC metrics exhibited cellular composition and gene expression profiles and low stress signatures, closely aligned with those of the developing human cerebral cortex. We developed CortiCOSI, a tool for assessing the quality of cortical organoids using scRNA-seq. Application of CortiCOSI to scRNA-seq datasets enabled the resolution of excitatory neuron phenotypes that were not detectable without rigorous QC stratification. This refined analysis identified reduced PP2A-mediated dephosphorylation and prefoldin complex dysfunction as potential drivers of the hyperphosphorylated and oligomeric tau phenotypes observed in *MAPT* V337M organoids.

Individual hPSC lines differ in their propensity for spontaneous differentiation and responsiveness to guided differentiation protocols.^45,46,69–74^ To accommodate this biological variability across large donor collections and CRISPR-edited lines, we optimized multiple stages of cortical organoid production. A key modification was the use of sustained-release native FGF2, which improved pluripotency, increased consistency across lines, and reduced the emergence of mesendodermal progeny. Interestingly, even low numbers of mesendodermal cells may impair cortical organoid differentiation, potentially through the secretion of signaling factors that counteract cortical patterning cues. Prior studies of cortical organoids also identified clusters of non-ectoderm derived cell types, even with directed (guided-to-cortex) methods.^4,7,14,44,75–77^ Hence, this is a common issue that can be alleviated by PSC culture with sustained-release FGF2.

While growth of iPSCs with sustained-release FGF2 reduced the incidence of mesendoderm contaminants in organoids, we still observed variable acquisition of cortical cell fates across lines. Line-specific differentiation propensity has been described across ectodermal, mesodermal, and endodermal lineages. Approaches to address this variability include screening hPSC expression profiles to select high-performing lines or empirically testing optimal patterning conditions for each line. ^69–72^ For example, small changes in activin/nodal and BMP factors are needed for efficient cardiac differentiation from different hPSC lines.^71^ Among the patterning factors we tested, tailoring the SB431542 was most effective at achieving high-quality cortical organoids. We found that hPSC lines with higher *ALK5/TGFBR1* expression required increased SB431542 dosing to achieve robust forebrain specification. Together with our finding that hPSC lines benefiting from FGF2DISC culture exhibit low endogenous FGF2 expression at the pluripotent stage, these results suggest that baseline hPSC transcriptional profiles could serve as a prescreening metric to guide line-specific optimization of culture and patterning conditions.

In addition to addressing line-to-line consistency, transitioning from a dish-based method to a 96 slit-well format improved organoid uniformity, growth dynamics, batch consistency, and production efficiency. The slit-well system eliminates the need to transfer developing organoids between culture formats for long-term growth. Organoid growth plateaus by 2-3 months, and we have successfully maintained organoids in this format for as long as a year (not shown). The slit-well format removes the requirement for manual cutting of fused organoids, thereby reducing the risk of contamination or mechanical damage. Further, our protocol does not require the use of animal products such as embedding in Matrigel, a mouse sarcoma cell product, or fetal bovine serum-supplemented media, or culture shaking at any stage, unlike other approaches,^3,78–80^ improving translational potential. Future improvements to the organoid model will include assessment at later timepoints when astrocytes emerge and electrophysiological activity increases, and incorporation of vascular cells and microglia, which are important contributors to neurodegenerative processes. As protocols for organoid maturation and aging continue to advance,^81,82^ these developments will further enhance physiological relevance. We envision that this cortical manufacturing platform could be used for generating organoids from pools of different hPSC lines^78,83,84^, however, growth with FGF2DISCs and selecting lines with similar requirements for SB431542 would be important for consistency in cortical patterning.

A key advance of this study is the development of an end-to-end manufacturing QC platform that enables the consistent generation of well-patterned cortical organoids, benchmarked by the quality of the resulting organoid product. By integrating stage-specific checkpoints, this approach supports the efficient production of well-patterned organoids at scale and enables rational adjustment of patterning conditions to accommodate variability across hPSC lines. Importantly, no single QC metric was sufficient to predict outcome; rather, combining orthogonal metrics provided a robust strategy for defining high-quality organoids. This platform enables a “fail-early” approach to exclude suboptimal cultures, thereby reducing resource use and experimental noise, while also providing insight into the causes of manufacturing failure to inform strategies for improving subsequent batches. Integration of these manufacturing metrics with transcriptomic validation through CortiCOSI further strengthens the link between production quality and biological authenticity, highlighting the importance of rigorous QC as a prerequisite for detecting subtle, genotype-driven phenotypes in human organoid systems.

The application of CortiCOSI to an organoid dataset generated by the dish-based method^18^ revealed that substantial variation can exist even within isogenic pairs produced without rigorous manufacturing QC, and demonstrated that CortiCOSI-guided stratification enhances the rigor of isogenic pair analyses. This refined analysis identified early molecular alterations that precede the detection of abnormal hyperphosphorylation and oligomeric V337M-tau species in the organoids. Building on these observations, we predict that reduced PP2A activity, together with diminished prefoldin complex function, synergistically drives the production of pathological tau species. In Alzheimer’s disease, marked downregulation of PP2A activity is closely associated with tau hyperphosphorylation,^63^ and restoring PP2A function has been proposed as a therapeutic strategy for AD and other neurodegenerative diseases.^85^ Prefoldin subunits *Pfdn5* and *Pfdn6* were recently identified in a *Drosophila* screen of hTau^V337M^ -induced ommatidial degeneration as strong genetic modifiers of hTau^V337M^ cytotoxicity. Overexpression of *Pfdn5* was shown to suppress tau aggregation, whereas reduced levels exacerbated tau-induced neurodegeneration.^86^ Bioinformatic analyses evaluating genetic risk and protein–protein interactomes show that 4R tau (the 4-repeat isoform that includes exon 10) and its pathogenic variants (V337M and P301L) display altered interactions with the prefoldin complex.^87^ Furthermore, *in vitro* studies of Huntingtin demonstrate that knockdown of prefoldin subunits (*PFDN2* or *PFDN5*) leads to accumulation of pathogenic soluble small oligomeric aggregates, underscoring the role of the prefoldin complex in early stages of protein aggregation.^67^ Notably, these gene expression changes we describe were not detectable when analyzing the full dataset, but became evident when focusing specifically on isogenic pairs with well-patterned and consistent CortiCOSI scores.

The cortical organoid manufacturing platform and the CortiCOSI tool we have developed establish a foundation for systematically dissecting tauopathy mechanisms over time through in-depth functional analysis and phenotyping of cortical organoids across large line sets, as well as evaluating targeted interventions.

## Acknowledgements

We are indebted to the Regenerative Research Foundation for support, the NSCI NeuraCell core facility for organoid production, technical support, and distribution to collaborating labs, and to Se-Jin Yoon and Sergiu Pasca for training on the organoid protocol that formed the foundation of this study. We thank Russell Ulbrich of ScientiaLux and TissueGnostics for his invaluable imaging expertise and support with TissueFAXS.

Schematics were created with BioRender.com (**Figures 1A, B, 2A, 4A, 6A**).

***hPSC lines*** *MAPT* hPSC lines (**Table S1**) were provided through the support of the Tau Consortium of the Rainwater Charitable Foundation and are available from https://www.neuralsci.org/tau.

Data collection and dissemination supported by the ARTFL-LEFFTDS Longitudinal Frontotemporal Lobar Degeneration (ALLFTD) Consortium (U19: AG063911, funded by the National Institute on Aging (NIA) and the National Institute of Neurological Diseases and Stroke) (NINDS) and the former ARTFL & LEFFTDS Consortia (ARTFL: U54 NS092089, funded by the NINDS and the National Center for Advancing Translational Sciences; LEFFTDS: U01 AG045390, funded by the NIA and the NINDS). The authors acknowledge the invaluable contributions of the study participants and families and the assistance of the support staff at each participating site. Lines are deposited at the National Centralized Repository for Alzheimer’s Disease and Related Dementias (NCRAD), supported by U24 AG21886, NIA.

## Author Contributions

Conceptualization, TB, ST, SL.

Methodology, TB, ST, SL.

Software, KRB, LQ, LD, FF, SB.

Formal Analysis, TB, KRB, LQ, MM, CAS, SB, LD, CP, JH.

Investigation - TB, KRB, SL, LQ, KW, AR, KT, SKG, SB, LMO, KL, RL, HL, RC, SJ, MM, HFBM, CAS, ST.

Resources - ST, CMK, SJH, AMG, DHG, FJT, BT.

Data Curation - TB, KRB, LQ, LD, CP, SB.

Writing - Original Draft, TB, ST.

Writing – Review and Editing, TB, ST, KRB, LQ, SB, FF, LD, SB, RC, SJ, BLA, DCB, MM, HFBM, CAS, MCS, SJH, CMK, DHG, AMG.

Supervision – ST, JS, AMG, DHG, CMK, SJH, FJT, BT.

Project Administration - TB, ST, SL.

Funding Acquisition - ST, MCS, CMK, SJH, AMG, DHG.

## Declaration of interests

TB, ST, SL, JS patent pending related to FGF2DISCs.

AMG serves on the SAB/SRB for Genentech and Muna Therapeutics.

SJH advises Proximity Therapeutics, Psy Therapeutics, Frequency Therapeutics, Souvien Therapeutics, Vesigen Therapeutics, Sensorium Therapeutics, 4M Therapeutics, Ilios Therapeutics, and Entheos Labs. SJH. has received speaking/consulting fees from Amgen, AstraZeneca, Biogen, Merck, Regenacy Pharmaceuticals, Syros Pharmaceuticals, Juvenescence Life, and sponsored research or gift funding from AstraZeneca, JW Pharmaceuticals, Lexicon Pharmaceuticals, Vesigen Therapeutics, Compass Pathways, Atai Life Sciences, and Stealth Biotherapeutics. None of SJH’s declared interests had a role in the design or content of this article.

FJT consults for Immunai Inc., CytoReason Ltd, Cellarity, BioTuring Inc., and Genbio.AI Inc. and has ownership in Dermagnostix GmbH and Cellarity.

The other authors declare no conflicts.

## Resource Availability

### Lead Contacts

Further information and requests for resources and reagents should be directed to and will be fulfilled by the Lead Contacts, Sally Temple and Taylor Bertucci.

