## Supplemental Methods for "Integrated Manufacturing Platform for Cortical Organoids Reveals Early Phosphatase and Prefoldin Dysregulation in *MAPT* V337M Neurons"

### Experimental Model and Subject Details

See Key Resource Table for antibody and reagent information.

### Cell lines

This study focused on a set of 17 hPSC lines from 8 different donors obtained from the Tau Consortium cell line collection (https://www.neuralsci.org/tau);[1] 51 additional lines were examined, 47 lines were tested for cortical organoid production, **Table S1**. All lines were negative for mycoplasma, and karyotypically normal by G-banded Karyotype (WiCell Research Institute, Inc.). The hPSCs were maintained in six-well plates coated with growth factor-reduced Matrigel or Cultrex at 37$^{o}$C and 5% $CO_2$. The cultures were fed either with daily mTeSR1 medium or with twice-weekly mTeSR1 medium plus an FGF2DISC. When hPSCs were cultured with FGF2DISCs, one FGF2DISC was added to one well of a 6-well plate with 2 mL of mTeSR1 medium for approximately a week. The medium, but not the FGF2DISC, was replaced every 2-3 days based on culture confluency. In either culture method, hPSC cultures were clump-passaged using ReLeSR about once a week. Cells were not allowed to grow past 80% confluency. To prepare for organoid production, hPSC colonies were single-cell passaged using Accutase or TrypLE Express.

### Method Details

**Organoid production:** Organoids were generated at the NeuraCell Core Facility (Neural Stem Cell Institute, NY, USA). When hPSC cultures reached 70–80% confluency, the medium was aspirated, and wells were rinsed twice with DMEM/F12. Subsequently, 1.5 mL of TrypLE Express or Accutase was added per well of a 6-well plate and incubated for 7–10 minutes at 37 °C and 5% $CO_2$ until cells detached. Gentle trituration using a 1000 µL pipette was performed to obtain a single-cell suspension. Cells were transferred to a 50 mL conical tube and washed twice with DMEM/F12. Cells were then counted manually using a hemocytometer and resuspended at $1 \times 10^6$ cells/mL in mTeSR1 supplemented with 10 µM ROCK inhibitor Y-27632 (Tocris). Organoids were generated in dishes following previously described methods.[2]

For organoid setup in a 96-slitwell plate (S-bio, MS9096SZ), 0.5–$1 \times 10^6$ cells in 1 mL were added to 9 mL of mTeSR1 containing 10 µM ROCK inhibitor. 100 µL of this suspension was dispensed into each well (5,000–10,000 cells per well), aiming to form spheroids 350–500 µm in diameter. Plates were incubated overnight at 37 °C and 5% $CO_2$ to allow spheroid formation. On the following day (day 0 of differentiation), 20 mL DMEM/F12 was added, and the plate was gently rocked to wash; this wash was repeated once. The medium was then replaced with 14 mL differentiation Medium A consisting of E6 supplemented with 2.5 µM dorsomorphin (DM), 10 or 20 µM SB431542, and 2.5 µM XAV939. Lyophilized small molecules were freshly reconstituted and aliquoted for each round of organoid production.

From days 2–5, plates were fed daily by gently aspirating ~14 mL of pooled medium and replacing it with freshly prepared Medium A, achieving ~65% medium exchange. On day 6, the medium was switched to neural medium (NM; Neurobasal-A supplemented with B-27 minus vitamin A, GlutaMAX, and Antibiotic-Antimycotic) containing 20 ng/mL EGF and 20 ng/mL FGF2 (Medium B). Medium B was changed daily for 10 days, then every other day (~3×/week) for 9 days with ~65% medium exchange.

On day 25, the medium was replaced with NM supplemented with 20 ng/mL BDNF and 20 ng/mL NT3 (Medium C), with ~65% medium exchange every other day (3×/week). From day

43 onward, organoids were maintained in NM without added growth factors, with medium changes every other day (3×/week) using 15–20 mL per dish (~75% exchange).

After 2 months, if more than 25% of organoids had been removed, the remaining organoids were transferred to low-bind or non-tissue-culture-treated 6-well plates (10–15 organoids per 4 mL of medium). These cultures were maintained with ~50% medium exchange 3×/week. Organoids were harvested for QC at day 20 and 2 months, and analyzed by qPCR and immunohistochemistry (IHC). Cortical markers were assessed: PAX6 and FOXG1 at day 20, and CTIP2/BCL11B and TBR1 at 2 months.

**Fixation and frozen sectioning of organoids:** Organoids were fixed using 4% paraformaldehyde at 4°C for 2 hours for 20-day timepoints or overnight for older organoids. They were then rinsed three times in PBS and allowed to sink in 30% sucrose in PBS overnight. The organoids were placed in cryotrays (Seal N Freeze) with OCT compound (Tissue Tek, 4583), snap frozen in a slurry of dry ice and isopropyl alcohol in a Seal N Freeze box and stored at -80°C. Organoids were cryostat-sectioned sequentially at 20 µm thickness using a Leica cryostat (model CM3050S). Sections were placed on microscope glass slides, dried overnight, and stored at -20°C for subsequent immunohistochemistry.

**Immunohistochemistry staining for organoid sections:** Slides were stored frozen at -20°C. Slides were thawed, brought to room temperature (RT), and lines drawn to partition the edges of sections on the glass slides using an ImmEdge Pen. The sections were rehydrated, blocked, permeabilized and immunostained with primary antibodies: PAX6 (1:10-1:100), FOXG1 (1:500), CTIP2 /BCL11B(1:500), SATB2 (1:500), BTUB III (1:1000), MAP2 (1:1000-2000),

TBR1 (1:500), GFAP (1:500). Primary antibodies were incubated overnight at 4$^{o}$C, then sections were washed three times with PBS and incubated with the corresponding Alexa Fluor conjugated secondary antibodies (1:333-1000) for 1 hour at RT. Sections were again washed three times, coverslipped, and imaged using fluorescence microscopy (Zeiss AXIO Observer.Z1; Zeiss 780; TissueFAXS (TissueGnostics)). See Key Resource Table for antibody information.

**Organoid Imaging with TissueFAXS:**

Fluorescence imaging of immunostained organoid sections was performed on the TissueGnostics TissueFAXS Q Confocal Tissue Cytometer, equipped with SpectraSplit 7 filters, Spectra X LED illumination and a Hamamatsu ORCA Fusion BT Camera, using widefield settings. Images were collected with a 20x air objective (NA 0.8), pixel size was maintained at 0.344 x 0.344 µm; total image size varied based on the number of tiles required to capture organoid sections of varying size.

**Organoid RNA isolation for qPCR:** Three organoids were collected randomly across the different quadrants of the 96 slit-well plate, then pooled. Total RNA was isolated from the organoids with Trizol RNA Clean and Concentrator Kit. RNA concentrations were measured by NanoDrop (Thermo Fisher), and cDNA was generated using the High-Capacity cDNA Reverse Transcription Kit. Real-time quantitative PCR was performed using Power SYBR Green Master Mix. GAPDH was used as the housekeeping gene. Ct values for GAPDH averaged approximately 20 ± 0.5 across all conditions.  The forward and reverse primers used are shown in the table below.

| qPCR Primers | | |
|---|---|---|
| **Primer** | **Forward Sequence** | **Reverse Sequence** |
| GAPDH | GAACGGGAAGCTTGTCATCAA | ATCGCCCCACTTGATTTTGG |
| PAX6 | CCCCACATATGCAGACACACA | GAACTGACACACCAGGGGAAA |
| FOXG1 | GCCAGCAGCACTTTGAGTTA | TGAGTCAACACGGAGCTGTA |
| TBR1 | ACGAACAACAAAGGAGCTTCA | TGGTACTTGTGCAAGGACTGTA |
| MAP2 | CAACGGAGAGCTGACCTCA | CTACAGCCTCAGCAGTGACTA |
| CTIP2/ BLC11B | CGAAAGGCATCTGTCCCAAG | ATGAGTGAGGGTGGGAGGA |
| COL1A2 | CCTGGTGCTAAAGGAGAAAGAGG | ATCACCACGACTTCCAGCAGGA |
| POU5F1 | CCTGAAGCAGAAGAGGATCACC | AAAGCGGCAGATGGTCGTTTGG |
| NANOG | CTCCAACATCCTGAACCTCAGC | CGTCACACCATTGCTATTCTTCG |
| SOX2 | GCTACAGCATGATGCAGGACCA | TCTGCGAGCTGGTCATGGAGTT |
| SOX17 | ACGCTTTCATGGTGTGGGCTAAG | GTCAGCGCCTTCCACGACTTG |

**Flow cytometry of hPSCs:** When hPSC cultures reached about 80% confluency, the medium was aspirated, and wells were rinsed twice with DMEM/F12. 1.5 mL of TrypLE Express or Accutase was added per well of a 6-well plate and incubated for 7-10 minutes at 37 $^{o}$C, 5% $CO_2$, monitoring by microscopy until cells detached from the dish. Using a 1000 µL pipette, gentle trituration was performed to achieve a single-cell suspension, which was then transferred to a 15 mL conical tube. Cells were washed twice with DMEM/F12. Following, the cells were resuspended in 1mL of PBS without calcium and magnesium (PBS-/-) and 9 mL of ice-cold methanol was slowly added while vortexing. On the day of staining, fixed cells were washed three times with PBS-/-. Cells were then counted using a hemocytometer and resuspended in 3% BSA/PBS-/- to a final concentration of 1 million cells per 100 µL. One million cells were incubated with TRA-1-60 (5 µL/ million cells), SSEA4 (20 µL / million cells), and SSEA1 (20 µL / 1 million cells) antibodies for 30 minutes at RT. After 2 washes with PBS-/- the stained cells were analyzed using a BD FacsAria II Flow Cytometer/Cell Sorter. Single-stained and unstained controls were used to set the gates (**see Figure S2D**). See Key Resource Table for antibody information.

**SDS-PAGE Western Blots:** For **Figures S1C,** organoids were randomly selected from the plate, then lysed in RIPA buffer supplemented with 1% HALT protease/phosphatase inhibitors 80 U/mL Benzonase, 10 mM DTT and 2% w/v SDS. Organoids were mechanically disrupted by vigorous trituration, incubated for 15 minutes at RT and 15 minutes on ice, followed by water bath sonication for 5 minutes, vortexing once halfway through. Lysates were cleared by centrifugation for 30 min at 20,000g at RT and supernatants were transferred to fresh Eppendorf tubes. Total protein concentrations were determined via BCA assay and 30 μg of protein per sample was separated using Tris-Acetate NuPAGE gel. Proteins were transferred to polyvinylidene difluoride (PVDF) membranes following standard wet transfer protocols. Membranes were blocked in 5% BSA in TBS-T (0.2% Tween) for 1 hour at RT and incubated with primary antibodies (pSer396 Tau (1:1000); Tau-5 (1:1000); betaIII-tubulin (1:10000); β-Actin (1:10000) diluted in blocking buffer overnight at 4 ºC. Following washes, the blots were incubated with corresponding secondary antibodies conjugated to HRP (1:10000) for 1.5 hours at RT, washed, activated with ECL reagent, and developed using standard darkroom techniques. Quantification was performed using Photoshop and ImageJ.

**Blue Native (BN)-PAGE Western Blots:** For **Figures 6D**, organoids were randomly selected and lysed using the NativePAGE Sample Prep Kit according to the manufacturer's protocol. Briefly, single organoids were homogenized in 90 μL of lysis buffer supplemented with 1% digitonin and 25 U/ml Benzonase by vigorous, repeated pipetting, then incubated on ice for 30 minutes. After incubation, the lysates were centrifuged at 20,000g for 30 minutes at 4 °C, and the supernatant was transferred to fresh centrifuge tubes. The protein concentration was determined by BCA, and the lysates were stored at -80°C. The samples were thawed, and G-250 from the sample prep kit was added fresh to a final concentration of 0.25% v/v (1/4 of the digitonin concentration). Equal amounts of total protein were loaded onto NativePAGE Novex 4-16% Bis-Tris Gels together with the NativeMark unstained protein ladder as size marker. Gels were run according to the manufacturer's recommendation with the NativePAGE Running Buffer Kit at 4°C and 150V constant for 60 minutes, first with the light blue cathode buffer. After 60 minutes, the light-blue cathode buffer was replaced with the dark-blue cathode buffer, the voltage was increased to 250V, and the gels were run for an additional 90 minutes. Proteins were transferred to PVDF membranes using standard wet transfer methods in NuPAGE transfer buffer at 200V at 4°C for 2 hours. After transfer, the membranes were destained in methanol to remove the blue G-250 dye and stained with Ponceau S. The size marker bands were marked with a pencil, and the membranes were destained again with water. From here on, the protocol followed the standard western blot procedure described for the non-native conditions for the data in **Figures S1C**.

**hPSC RNA extraction and RNA-seq library preparation (Figure 2):** RNA was extracted from hPSC samples using the Zymo Research Quick-RNA Miniprep Plus kit. The concentration was measured with the Qubit RNA High Sensitivity Assay kit. The RNA Integrity Number was evaluated for each sample on an Agilent TapeStation (all RIN > 9). Library preparation was completed using the Illumina TruSeq Stranded Total RNA kit. 100 bp paired-end sequencing was completed on an Illumina NovaSeq 6000 to acquire >50 million fragments per sample. For

FGF2DISC vs daily-fed samples from the same hPSC line, RNA extraction and library construction were performed on the same day by the same operator to reduce batch effects.

**hPSC RNA-seq preprocessing and differential expression analysis (Figure 2):** Adaptor-trimmed RNA sequencing files were mapped to the genome (GRCh38; Homo_sapiens.GRCh38.dna.primary_assembly.fa; https://ftp.ensembl.org/pub/release-106/fasta/homo_sapiens/dna/using RNA-STAR[3] (v2.7.10a). Gencode annotations (Release 40 (GRCh38.p13)) were used from ENSEMBL. Samples sequenced on different lanes were merged using Samtools (v1.15). Finally, RSEM[4] (v1.3.3) was used for quantification to obtain expected read counts at the gene and transcript levels.

Expected gene read counts were then subjected to several processing steps in preparation for downstream analysis, primarily in R. First, counts per million (CPM) were computed from gene-level counts for filtering purposes. Genes were first filtered using the filterByExpr() function in edgeR[5] (v4.0.16) using the default settings. Genes with an effective length (measured by RSEM) of less than 15 bp were also removed. The counts for the remaining genes (19,570) passing these filters were normalized using edgeR with an adjustment for sample read depth variance. An offset value calculated with CQN[6] accounting for GC content bias and gene effective length bias in read quantification was also incorporated during the normalization process.

To identify differentially expressed genes between hPSCs grown with FGF2DISCs versus daily feeding, a linear mixed-effect model (LMM) was used to account for variations across different iPSC clones with variancePartition[7] (v1.32.5) R package. RUVSeq[8] (v1.36.0) was used to account for unknown variance not explained by medium conditions or lines. A final

LMM model was determined to be ~Condition+(1|line)+W_1+W_2+W_3+W_4, where Condition refers to culture medium conditions (FGF2DISC vs. daily feeding), (1|line) is a random effects variable for the name of the iPSC lines and W_1 to W_4 are factors of unwanted variation estimated by RUVr. The number of k in RUVr was selected based on sample size. Subsequently, voomWithDreamWeights(), dream() with Kenward-Roger approximation and eBayes() functions were used to extract genes significantly different between FGF2DISC and daily feeding hPSC culture conditions with FDR < 0.05 and absolute log2 fold change (logFC) > 0.3.

**hPSC RNA-seq Gene Ontology analysis (Figure 2D, Supplemental Table S2):** g:ProfileR2[9] was used for gene ontology enrichment. 'BP' (biological process) terms, KEGG pathways and Reactome pathways were included in the analysis. Results were first filtered for minimum term size of 30 and FDR-adjustment for p-values. The significant terms were clustered by first computing a similarity matrix from pairwise overlap coefficients, which were then converted into a distance matrix. Hierarchical clustering was used on the resulting distance matrix. Each cluster was marked with the term with the lowest FDR.

**hPSC RNA-seq correlation and distance analysis across samples (Figure S2A):** We calculated Spearman's correlation and Euclidean distance across all possible pairs of samples within FGF2DISC and daily feeding conditions using raw logCPM. Fisher's Z-transformation was applied to the Spearman's rho values, which were used for a paired Student's t-test between FGF2DISC and daily feeding groups. Paired Wilcoxon signed-rank test was applied to compare distances between FGF2DISC and daily feeding groups. The standard deviations of raw logCPM of all expressed genes/consensus peaks were calculated within FGF2DISC or daily feeding groups. Between group comparisons were done using the paired Wilcoxon-signed rank test.

**Slit-well organoid dissociation and cell hashing for scRNA-seq (Figures 3-6):** Three organoids were collected randomly across the different quadrants of the 96 slit-well plate and pooled. Organoids were incubated in Papain for 90-120 minutes at 37°C with gentle shaking (300 rpm) and regular pipetting with a glass Pasteur pipette to obtain single cell suspensions. Cells were pelleted and resuspended in ice-cold PBS before being passed through a 40 µm filter and kept on ice. Viability and cell number was determined by Trypan Blue staining and counting using the Countess system. Three organoids per condition, totaling 150-300,000 cells, were submitted to the New York Genome Center (NYGC) for hashtag-oligonucleotide (HTO) antibody binding, library preparation, and sequencing as previously described[10,11].

**Slit-well organoid scRNA-seq data QC, alignment, and cell annotation:** Data QC and alignment were carried out at the NYGC. Briefly, analysis up to the counting of UMIs per cell per gene was carried out using Drop-Seq tools (Drop-seq tools v1.0, McCarroll Lab), and reads were aligned to the human hg38 genome using the STAR aligner[3]. HTOs were identified and counted using Cite-Seq Count (https://hoohm.github.io/CITE-seq-Count/), and the samples were de-multiplexed using HTODemux.[12] Following processing at NYGC, the data was further processed using Seurat 3.0 dev[12,13] to filter out cells with < 200 detectable genes, < 500 reads and > 20% mitochondrial rate. Cells were initially annotated using the SingleR package,[14] using custom reference libraries constructed from human fetal neocortical single-cell RNA sequencing data[15] and bulk adult astrocyte RNA-sequencing data.[16] The top 3,000 most variable genes were used as input for cell type annotation. Loupe Browser software was used to visualize overall composition overlap between hPSC donors and SB431452 levels (see **Figures 3C**). In order to annotate off-target non-neural cell clusters, cells were again annotated using the SingleR package, but with the human cell atlas of fetal gene expression[17] as reference data (**see Figure S3**).

**Glycolysis Metabolic Stress Assessment (Figure 5):** Metabolic stress scores were computed as described in the Human Neural Organoid Cell Atlas (HNOCA) publication.[18] Briefly, the three primary fetal brain datasets used for the analogous comparison in the HNOCA publication,[19-21] together with the HNOCA dataset, were subset to dorsal telencephalic neurons. Following CPM normalization and log1p-transformation of counts, cells with fewer than 200 expressed genes were removed from each dataset. Cells that originated from a study that contributed fewer than 20 cells to the subset HNOCA dataset at this point were also removed. Datasets were concatenated using the intersection of the genes reported across all datasets. The Molecular Signatures Database gene set "hallmark glycolysis" [22,23] was used as a proxy for metabolic stress. Of the 200 genes in the gene set, 192 were reported across all datasets in this comparison and hence used for scoring (**Supplementary Table S7**). The scanpy (81) (v.1.9.3) function score_genes was then used to compute a stress score per cell.

**Gruffi Stress Assessment:** Cellular stress associated with organoid patterning was assessed using Gruffi v1.5.6[24] with default parameters. Gruffi is a computational method for detecting and quantifying cellular stress signatures in single-cell RNA sequencing (scRNA-seq) data. It identifies stressed cells based on the expression of defined stress-response gene programs and assigns a stress score to each cell, enabling classification of cells by stress state**.** scRNA-seq data were analyzed to identify the percentage of stressed neurons (ExDp1, ExDp2, ExM, ExMU, ExN, CGE, MGE and UN). These values were compared at the sample level to evaluate relationships between stress signatures and patterning outcomes (**Figure 5B**).

**Regional Transcriptomic Quality Assessment Using VoxHunt and Generation of the Cortical Regional (CR) Identity Value**: To assess organoid regional identity and transcriptomic fidelity relative to expected cortical developmental trajectories, scRNA-seq datasets were analyzed using the VoxHunt framework. VoxHunt compares single-cell transcriptomic profiles to reference developmental brain atlases, enabling quantification of similarity between experimental cells and anatomically defined in vivo brain regions.

For each organoid sample, transcriptomic correlations were evaluated either within the annotated excitatory neuron subset or across all cells in aggregate. These comparisons were performed against curated cortical developmental reference datasets, including the Allen Developing Mouse Brain Atlas and human BrainSpan datasets, with a focus on dorsal forebrain/neocortical identity.

The cortical regional (CR) identity value, calculated across all cells, was used as a transcriptomic quality control metric to assess organoid regional specification and developmental consistency across samples. The CR value was defined as the ratio of dorsal pallium (mouse) or neocortex (human) correlation to the mean correlation across all other brain regions. Samples showing low similarity to expected cortical reference regions or increased correlation with off-target developmental identities were classified as lower-quality organoids. CR values were derived using both human and mouse developmental reference atlases to ensure robustness in regional identity assignment. BrainSpan represents a human dataset but is smaller than the ABA mouse reference. As larger and more comprehensive human developmental reference datasets become available, the CortiCOSI scoring framework can be readily adapted to incorporate them.

**CortiCOSI Scores (Figure 4F, S5, S8)**: To assess organoid quality holistically, we combined the CR identity values and gene expression levels into a single metric. For each organoid sample, individual metrics were first evaluated independently. The CR value was then integrated with the expression levels of *FOXG1* and *COL1A2*, two genes selected for their relevance to the quality and consistency of cortical organoid production. Thresholds were established for each individual metric to define well-patterned samples. These three metrics had limited discriminatory power when used independently. Thus, these metrics were combined into a composite z-score, enabling quantification of inter-sample variation and robust discrimination between pass and fail quality control groups with minimal overlap. Although z-scores are relative measures, integrating these three orthogonal benchmarks into a single metric (CortiCOSI) enabled visualization that can be applied to other datasets to assess inter-sample organoid consistency across the scRNA-seq dataset.

**Reanalysis of scRNA-seq V337M dataset (Figure 6)**: Single-cell RNA sequencing (scRNA-seq) analyses were performed in three major stages: (1) transcriptomic quality assessment of cortical organoids using CortiCOSI scoring, (2) preprocessing and normalization of filtered datasets for downstream analysis, and (3) differential gene expression (DGE) analysis using a pairwise meta-analytic framework.

Following CortiCOSI assessment, samples were identified for CortiCOSI-selected downstream analysis. Single-cell gene expression matrices were processed using the Seurat framework. Cells were filtered according to established QC criteria, including removal of low-quality cells and off-target populations identified through annotation and transcriptomic profiling.

Gene expression counts were log-normalized to account for differences in sequencing depth across cells. Highly variable genes were identified for downstream scaling and dimensionality reduction. Expression values were scaled to center and standardize gene expression across cells while regressing technical sources of variation where appropriate. Cell type annotations were used to subset relevant populations for downstream DGE analysis, with analyses focused on excitatory neuronal populations using log-normalized TBR1>2 expression levels as a filter.

DGE analysis was performed using the scMetaIntegrator framework, which implements a meta-analytic strategy optimized for single-cell datasets with paired experimental designs. Given the isogenic paired structure of the dataset, differential expression was first computed independently within each matched pair rather than pooling all cells into a conventional case-control framework, thereby reducing pseudoreplication bias and preserving pair-specific biological variability.

Pairwise differential expression results were subsequently integrated using scMetaIntegrator's meta-analysis framework to identify genes exhibiting consistent genotype-associated expression changes across pairs. Effect size estimates, heterogeneity metrics, and combined statistical significance measures were generated for each gene. Primary differentially expressed gene (DEG) prioritization was based on effect size-derived statistical measures, which provide a more conservative estimate of genes showing reproducible differential expression across matched comparisons.

Visualization of DGE results, including volcano plots, forest plots, and pairwise effect summaries, was performed using the accompanying scMetaIntegrator visualization tools, custom R scripts and GraphPad Prism.

**Statistical Strategy for Differential Gene Expression Identification:**

Power calculations performed using study simulation tools indicated potential limitations in distinguishing distinct gene expression distribution profiles when identifying DEGs.[25] Consequently, parametric testing approaches were excluded due to the elevated risk of Type I error associated with applying these methods to non-normally distributed data.[26] Furthermore, the use of paired isogenic samples for DEG analysis has been shown to enhance both sensitivity and specificity, whereas unpaired analyses are more susceptible to technical variability, increasing the likelihood of false-positive findings.[27] Although pseudobulk strategies are commonly employed for DEG testing in single-cell RNA sequencing data, they present notable limitations in specific contexts, particularly when compared to approaches that leverage pairwise designs.[28]

**Statistics:** All statistical analysis and data visualization was performed with GraphPad Prism 10.4.1, unless otherwise stated. Data are presented as mean ± standard deviation unless otherwise indicated. Distribution of the raw data was tested for normality of distribution using the Shapiro-Wilk test and the Kolmogorov-Smirnov test. If normally distributed, statistical analyses were performed using the unpaired two-tailed t-test or one-way ANOVA with the Tukey test post hoc unless otherwise stated. If the data were not normally distributed, nonparametric tests were used (e.g., the Mann–Whitney U test for unpaired comparisons or Spearman's rank correlation for association analyses). Box plots were created using GraphPad. The style selected was entitled "Box and whiskers" with the option for whiskers extended to the minimum and maximum values with all points shown. The center line represents the median; box limits are to the $25^{th}$ to the $75^{th}$ percentile. Violin plots include lines to indicate quartiles of the respective distributions.

| Key Resource Table | | | | |
|---|---|---|---|---|
| **Antibodies** | | | | |
| **Reagent** | **Example Company Validation data*** | **Example References using this antibody** | **Source** | **Identifier (Lot)** |
| Tra-1-60 | Flow cytometry on hPSCs | [29,30] | BD Bioscience | 561153 (3279907) |
| SSEA4 | Flow cytometry on hPSCs | [31] | BD Bioscience | 560796 (3016392) |
| SSEA1 | Flow cytometry of E14 mouse PSCs | [32] | BD Bioscience | 560127 (3173967) |
| PAX6 | WB and flow data | [33] | DSHB | PAX6 antibody registry ID: AB_528427 |
| FOXG1/BF1 | IHC | [34] | Takara | M227 (AJ12076A, AK21911A) |
| CTIP2/BCL11B | WB, Flow and ICC | [35] | Abcam | ab18465 (GR32772266-5, GR3272266-22) |
| TBR1 | WB, Flow and IHC | [36] | Abcam | ab31940 (GR3217079-1, GR3371747-1) |
| SATB2&1 | WB, Flow and ICC | [37] | Abcam | ab51502 (GR3235758-5, GR3381222-4) |
| GFAP | ICC and IHC | [38] | Aves Labs | AB_2313547 Lot# GFAP857944 |
| betaIIItubulin | WB, ICC, IHC | [39] | Sigma | T-8660 (097M4835V, 0000091060, 00000116046, 00000139791) |
| MAP2 | WB, ICC, IHC | [40] | SYSY (Synaptic Systems) | 188004 (4-38, 3-32, 3-34, 5-40) |
| goat anti ms IgG1 ALEXA488 | Lot-specific QC available; specificity determined by antigen/binding assay or ELISA | [41] | Jackson | 115-545-205 (150562, 153248, 162833) |
| goat anti rabbit Cy3 | Lot-specific QC available; specificity determined by immune-electrophoresis or ELISA | [42] | Jackson | 111-165-144 (140782, 147422, 150960) |

| | | | | |
|---|---|---|---|---|
| goat anti rat ALEXA-488 | ICC | [43] | Invitrogen | A-11006 (2161035, 2299157, 2480078) |
| goat anti ms IgG1 ALEXA546 | ICC; Spectral data | [44] | Invitrogen | A-21123 (228626, 2438513, 2132524) |
| goat anti ms IgG2b ALEXA647 | | | Invitrogen | A-21242 (1968702, 2379469) |
| goat anti gp ALEXA-647 | Lot-specific QC available; specificity determined by immunoelectrophoresis or ELISA | [45] | Jackson | 106-605-003 (153308, 160728) |
| Total Tau-5 | WB, IHC | [46] | Biolegend | 806401 (B350881) |
| p-Tau-Ser396/Ser404 (PHF1) | WB, IHC | [47] | gift from Peter Davies, availability and validation data https://www.alzforum.org/alzantibodies/tau-phf-1; contact | |
| 4R-Tau (ET3) | WB, ICC, IHC | [48] | gift from Peter Davies, for availability contact | |
| GAPDH | WB, ICC | [49] | Abcam | ab8245 |
| HRP anti mouse | CoA available | [50] | Cell Signaling | 7076 (L38) |
| HRP anti rabbit | CoA available | [51] | Cell Signaling | 7074 (L33) |
| HRP goat anti guinea pig | WB, ELISA, immunoelectrophoresis | [52] | Invitrogen | A18769 (93-37-072222) |
| pSer396 Tau | ICC, WB | [53] | Thermo Fisher Scientific | 44-752G (2302647) |
| Tau-5 | WB, ICC, IHC | [54] | Thermo Fisher Scientific | AHB0042 (VA293183) |
| betaIII tubulin | WB, ICC, IHC | [55] [56] | SYSY (Synaptic Systems) | 302304 (302304/1-8) |
| β-actin | WB, ICC, IHC | [57] | Sigma | A1978 (0000137631) |
| GAPDH | WB, ICC, IHC | [58] | Cell Signaling | 2118 (gr3316865-13) |

*In lab antibody validation: IHC/ICC:* Primary antibodies are screened at different concentrations by immunostaining to determine the optimal concentration and that the staining location and distribution are as expected; tissue/cells known to express the marker are used as positive controls; negative controls include secondary only controls, pre-immune serum when available, and knockout lines when available; controls are performed at the same time as experimental staining. *WB:* Antibodies are tested to confirm that they label bands of the expected size, with appropriate positive and negative controls as described. *Bridging studies:* When a new lot is procured, it is tested against the prior lot to ensure consistency.

| **Chemicals, peptides, and recombinant proteins** | | |
|---|---|---|
| **Reagent** | **Source** | **Identifier** |
| DMEM/F12 | Corning | 10-092-CV |
| Essential 6 medium (E6) | Life Technologies | A1516401 |

| | | |
|---|---|---|
| Essential 8 medium (E8) | Life Technologies | A1517001 |
| mTeSR1 medium | StemCell Tech | 85850 |
| FGF2DISCs | StemCultures LLC | DSC500 |
| Neurobasal-A (- L-glutamine) | Life Technologies | 10888-022 |
| Accutase | Innovative Cell Tech | AT-104 |
| TrypLE Express | Gibco | 12605010 |
| Antibiotic-Antimycotic (Anti A) 100X | Thermo Fisher Scientific | 15240-062 |
| B-27 supplement without vitamin A | Thermo Fisher Scientific | 12587001 |
| GlutaMAX Supplement | Thermo Fisher Scientific | 35050061 |
| Matrigel | Thermo Fisher Scientific | 356230 |
| Cultrex | Thermo Fisher Scientific | 343400502 |
| Anti-Adherence Rinsing Solution | StemCell Tech | 7010 |
| Dorsomorphin | Tocris | 3093 |
| SB431542 | Tocris | 1614 |
| XAV-939 | Tocris | 3748 |
| IWR-endo | Tocris | 3532 |
| LDN | Tocris | 6053 |
| Y-27632 | Tocris | 1254 |
| human BDNF | Peprotech | 450-02 |
| human BDNF | Shenandoah Biotech. | 100-01 |
| human EGF | Peprotech | AF-100-15 |
| human EGF | Shenandoah Bioteh. | 100-26 |
| human NT3 | Peprotech | 450-03 |
| human NT3 | Shenandoah Biotech. | 800-06 |
| rhFGF2 | Peprotech | 100-18B |
| rhFGF2 | Shenandoah Biotech. | 100-28 |
| Trizol | Thermo Fisher Scientific | 15596018 |
| RNA Clean and Concentrator Kit | Zymo Research | R1014 |
| High-Capacity cDNA Reverse Transcription Kit | Thermo Fisher Scientific | 4368814 |
| SYBR Green Master Mix | Thermo Fisher Scientific | 4368702 |
| Papain Dissociation System | Worthington | LK003150 |
| RIPA Buffer | Thermo Fisher Scientific | 89900 |
| Protease inhibitor cocktail | Roche | 11697498001 |
| Phosphatase inhibitor cocktail | Roche | 4906845001 |
| DC Assay | Bio-Rad | 5000111 |
| Laemmli SDS sample reducing buffer | Boston Bioproducts | BP-111R |
| PVDF Transfer Kit | Bio-Rad | 1704272 |
| BSA | Fisher Scientific | BP9703100 |
| TBS (10X) | Bio-Rad | 1706435 |

| | | |
|---|---|---|
| Tween-20 | MP Biomedicals | 11TWEEN201-CF |
| ECL substrate | Bio-Rad | 1705061 |
| Quick-RNA Miniprep Plus kit | Zymo Research | R1057 |
| Qubit RNA High Sensitivity Assay kit | Thermo Fisher Scientific | Q32852 |
| High Sensitivity RNA ScreenTape | Agilent | 5067-5579 |
| TruSeq Stranded Total RNA Library Prep Gold | Illumina | 20020598 |
| IPA Buffer | Boston BioProducts | BP-115-500 |
| HALT protease/phosphatase inhibitor | Thermo Fisher Scientific | 78444 |
| Benzonase Nuclease | Sigma-Aldrich | E1014 |
| DTT | New England BioLabs | B7705SVIAL |
| Laemmli SDS sample reducing buffer | New England BioLabs | B7703S |
| PVDF membranes | Millipore Sigma | IPVH00010 |
| BSA | Sigma-Aldrich | A7906 |
| Sodium dodecyl sulfate | Sigma-Aldrich | 71725 |
| TBST | Boston BioProducts | IBB-180 |
| Pierce™ ECL Western Blotting Substrate | Thermo Fisher Scientific | 32106 |
| SuperSignal™ West Pico PLUS ECL substrate | Thermo Fisher Scientific | 34580 |
| SuperSignal™ West Dura ECL substrate | Thermo Fisher Scientific | 34076 |
| BCA Assay | Thermo Fisher Scientific | 23227 |
| NuPAGE™ 7%, Tris-Acetate Mini Protein Gels | Invitrogen | EA03585BOX |
| NuPAGE™ Tris-Acetate SDS Running Buffer | Thermo Fisher Scientific | LA0041 |
| NativePAGE Sample Prep Kit | Thermo Fisher Scientific | BN2008 |
| NativePAGE Novex 4-16% Bis-Tris Gels | Thermo Fisher Scientific | BN1002BOX |
| NativeMark Unstained Protein Standard | Thermo Fisher Scientific | LC0725 |
| NativePAGE Running Buffer Kit | Thermo Fisher Scientific | BN2007 |
| NuPAGE transfer buffer | Invitrogen | NP00061 |
| Transfer buffer | Boston BioProducts | BP-190 |
| **Experimental Models: Cell Lines** | | |
| Refer to Table S1. | | |
| **Deposited Data** | | |
| hPSC RNAseq data will be deposited at GEO | GSE239811 | Token: ofutmqsabzgfhyh |
| Organoid scRNAseq data will be deposited at GEO | GSE239618 | Token: mdmtkckgfjofrul |

| Key Resource Table (cont.) | | |
|---|---|---|
| **Software and algorithms** | | |
| **Name** | **Reference** | **Link** |
| Image Lab software | n/a | https://www.bio-rad.com/en-us/product/image-lab-software?ID=KRE6P5E8Z |
| Adobe Photoshop v. 22.1.1 | Adobe | www.adobe.com/ |
| Image Studio Lite v. 5.2.5 | LI-COR Biosciences | https://www.licor.com/bio/image-studio/ |
| Samtools (v1.15) | n/a | http://www.htslib.org/ |
| Picard tools (v2.5.0) | n/a | https://broadinstitute.github.io/picard/ |
| RSEM (v1.3.3) | 4 | https://github.com/deweylab/RSEM |
| edgeR (v3.40.2) | 5 | https://github.com/StoreyLab/edge |
| CQN (v1.44.0) | 6 | https://bioconductor.org/packages/release/bioc/html/cqn.html |
| variancePartition (v1.28.9) | 7 | https://github.com/GabrielHoffman/variancePartition |
| g:ProfileR2 (v0.2.2) | 9 | https://github.com/egonw/r-gprofiler2 |
| Rstudio | n/a | https://posit.co/products/open-source/rstudio/ |
| ImageJ | n/a | ImageJ (nih.gov) |
| Seurat 3.0 (dev) | 59 | https://satijalab.org/seurat/ |
| SingleR | 60 | Using SingleR to annotate single-cell RNA-seq data (bioconductor.org) |
| VoxHunt | 61 | https://github.com/quadbio/VoxHunt |
| Gruffi | 24 | https://github.com/jn-goe/gruffi |
| Inkscape | n/a | https://inkscape.org/ |
| Loupe Browser 8.1.2 | n/a | https://www.10xgenomics.com/support/software/loupe-browser/latest |
| Gruffi | 24 | https://github.com/jn-goe/gruffi |

**Supplemental Tables**

**Table S1.** hPSC line information.

**Table S2.** Pathway enrichment analysis of hPSCs cultured with FGF2DISCs and fed twice weekly with mTeSR1, compared to hPSCs fed daily with mTeSR1 (see **Figure 2**).

**Table S3.** Differentially expressed genes in hPSCs cultured with FGF2DISCs and fed twice weekly with mTeSR1 compared to hPSCs fed daily with mTeSR1 (see **Figure 2**).

**Table S4.** Differentially expressed genes in off-target cell types (clusters 6, 14, 16, 17, and 19) compared to other clusters (see **Figure S3D**).

**Table S5.** Gene expression counts (log counts per million, CPM) per sample, used to compare hPSC lines and evaluate those that were unsuccessful under the organoid patterning conditions (10 μM SB431542) (see **Figure S4G**).

**Table S6.** scRNA-seq samples and associated metadata (see **Figures 3–5**).

**Table S7.** Hallmark glycolysis genes used in the glycolysis stress gene score shown in **Figure 5A**.

**Table S8.** HNOCA dataset samples corresponding to graphs shown in **Figure 5A**.

**Table S9.** Differentially expressed genes comparing CortiCOSI-selected V337M mutants versus corrected isogenic controls in excitatory neurons ($TBR1^{+}$ cells) (see **Figure 6**).

**Table S10.** Differentially expressed genes comparing all V337M mutants versus corrected isogenic controls in excitatory neurons ($TBR1^{+}$ cells) (see **Figure 6**).

**Ethics Statement**

The experiments utilizing human stem cells complied with all relevant guidelines and regulations, including guidelines issued by the International Society for Stem Cell Research (ISSCR) and NIH. All the lines used in the study were human induced pluripotent stem cells (hiPSCs); details regarding cell line sources, institutions responsible for oversight and review boards, and associated publications are provided in Table S1. Donors provided informed consent for the use of their cells to generate hiPSCs and derivative cell types for research purposes. This study employed cerebral cortical organoid technology, and development of the model was informed by ongoing ethical considerations surrounding neural organoid research.[62] The use of human stem cell-derived models enables investigation of disorders affecting the human cerebral cortex, including frontotemporal dementia (FTD), in a human-relevant context. The authors are committed to transparency and data sharing, and information regarding data and code availability is provided in the main manuscript.
