## Supplemental Figures for "Integrated Manufacturing Platform for Cortical Organoids Reveals Early Phosphatase and Prefoldin Dysregulation in *MAPT* V337M Neurons"

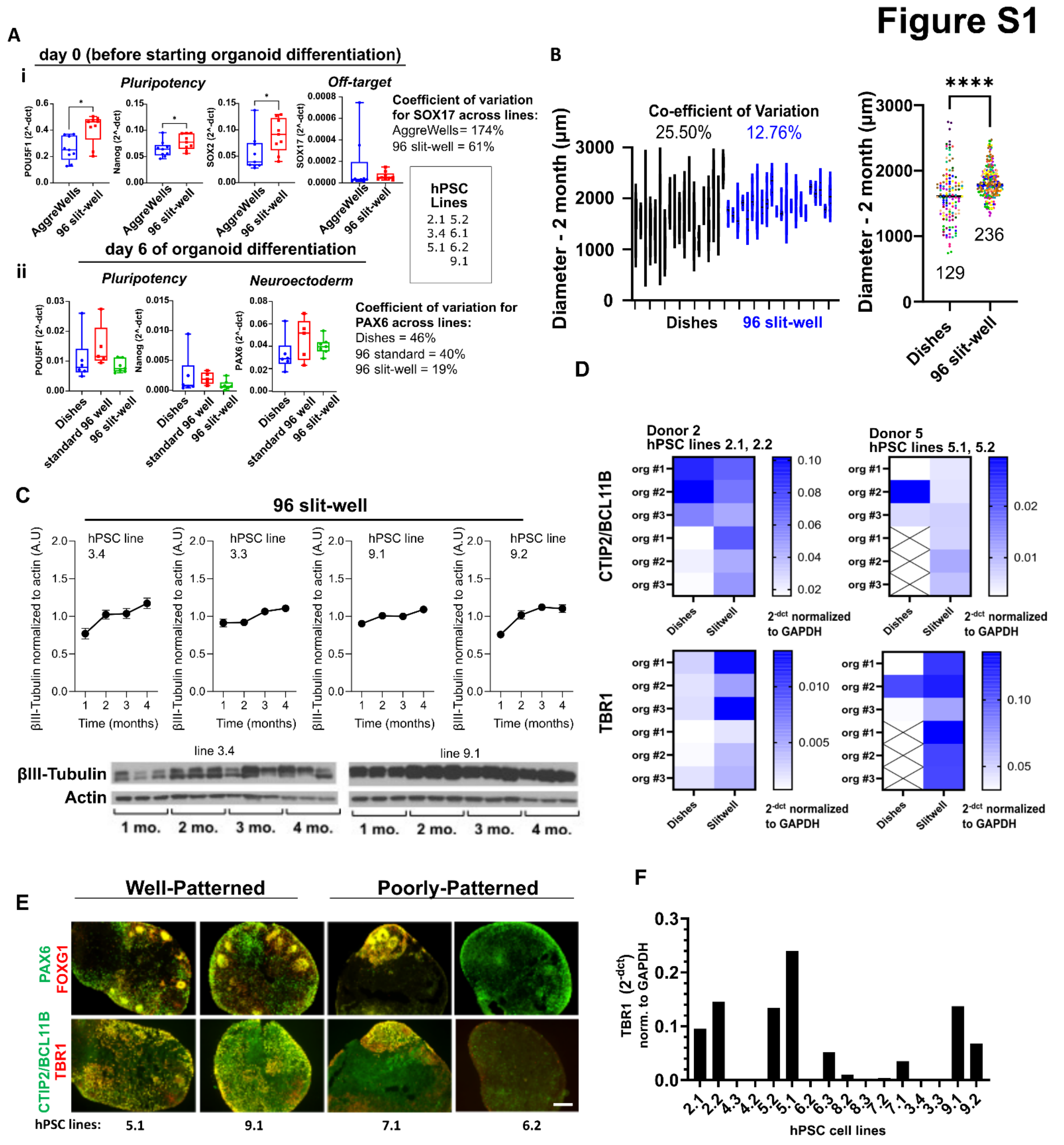


**Supplemental Figure 1. Characterization of cortical organoids across culture platforms.**

**S1A**) qPCR analysis (2^dCt normalized to *GAPDH*) of: i) Pluripotency markers and off-target *SOX17* in day 0 hPSC spheroids (AggreWells vs. 96-slitwell plates; n=9 samples/condition from 7 lines; Wilcoxon paired rank test). ii) *POU5F1, NANOG,* and *PAX6* in day 6 organoids (dishes vs. 96-well vs. 96-slitwell plates; n=7 lines).

Coefficients of variation (CV) reported for *SOX17* and *PAX6.* **S1B)** Diameter of 2-month-old organoids (dishes vs. 96-slitwell plates). Left: Range of 3–12 organoids per line/experiment (Dishes: n=19 samples from 15 lines; 96-slitwell: n=22 samples from 14 lines). Right: Individual organoid diameters colored by line (Dishes: n=129; 96-slitwell: n=236; two-tailed Mann-Whitney U test, $p < 0.0001$). **S1C)** Western blots and densitometry for βIII-tubulin (relative to Actin) over time (3 pooled organoids/timepoint). See **Fig. S9A** for raw blots. **S1D)** qPCR of deep-layer markers *TBR1* and *CTIP2/BCL11B* in individual 2-month-old organoids (n=12 samples from 4 lines). 'X' denotes insufficient material (<1% organoid efficiency). **S1E)** Representative immunofluorescent images of 2-month-old organoids stained for forebrain progenitor markers (PAX6, FOXG1) and deep-layer neuron markers (CTIP2/BCL11B, TBR1), n=4 lines. Scale bar = 200 μm. **S1F)** qPCR screen of 2-month-old organoids across 16 lines cultured in 96-slitwell plates (3 pooled organoids/column). Note inter-line expression variability.

### Figure S2

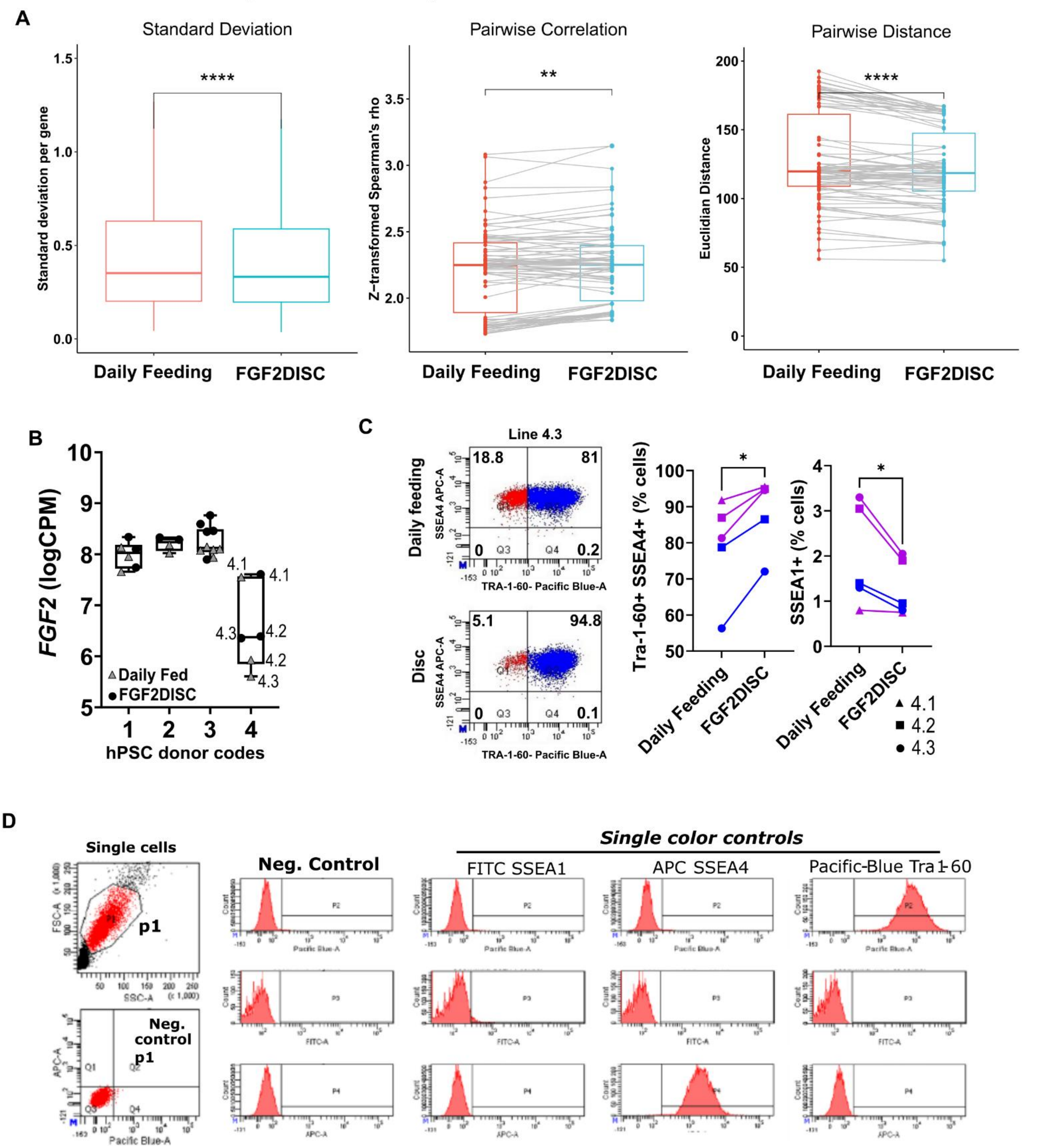


**Supplemental Figure 2. Impact on hPSC pluripotency following FGF2DISC culture.**

**S2A)** Variance across hPSC cultures computed from RNA-seq raw log CPM using SD (left), spearman's rho (middle) and Euclidean distance (right), comparing daily mTeSR1 feeding vs FGF2DISC + twice-weekly mTeSR1 (11 lines, 4 donors, 3 independent experiments; line 1.1 was included in each experiment as a control). **S2B)** *FGF2* expression across donors; culture method indicated by symbol shape (note lower expression of FGF2 in donor #4). Note that CRISPR-edited lines (4.2 and 4.3) show the lower *FGF2* expression levels. **S2C)** Donor #4 lines: flow cytometry of pluripotency markers TRA-1-60 and SSEA4 and differentiation marker SSEA1, comparing daily mTeSR1 vs FGF2DISC; line indicated by shape; 2–6 wells pooled/sample (5 samples/condition); experiments indicated by color (2 independent experiments); paired two-tailed t-test, *$p<0.05$. **S2D)** Flow cytometry gating for singlets and single-color controls.

**Figure S3**

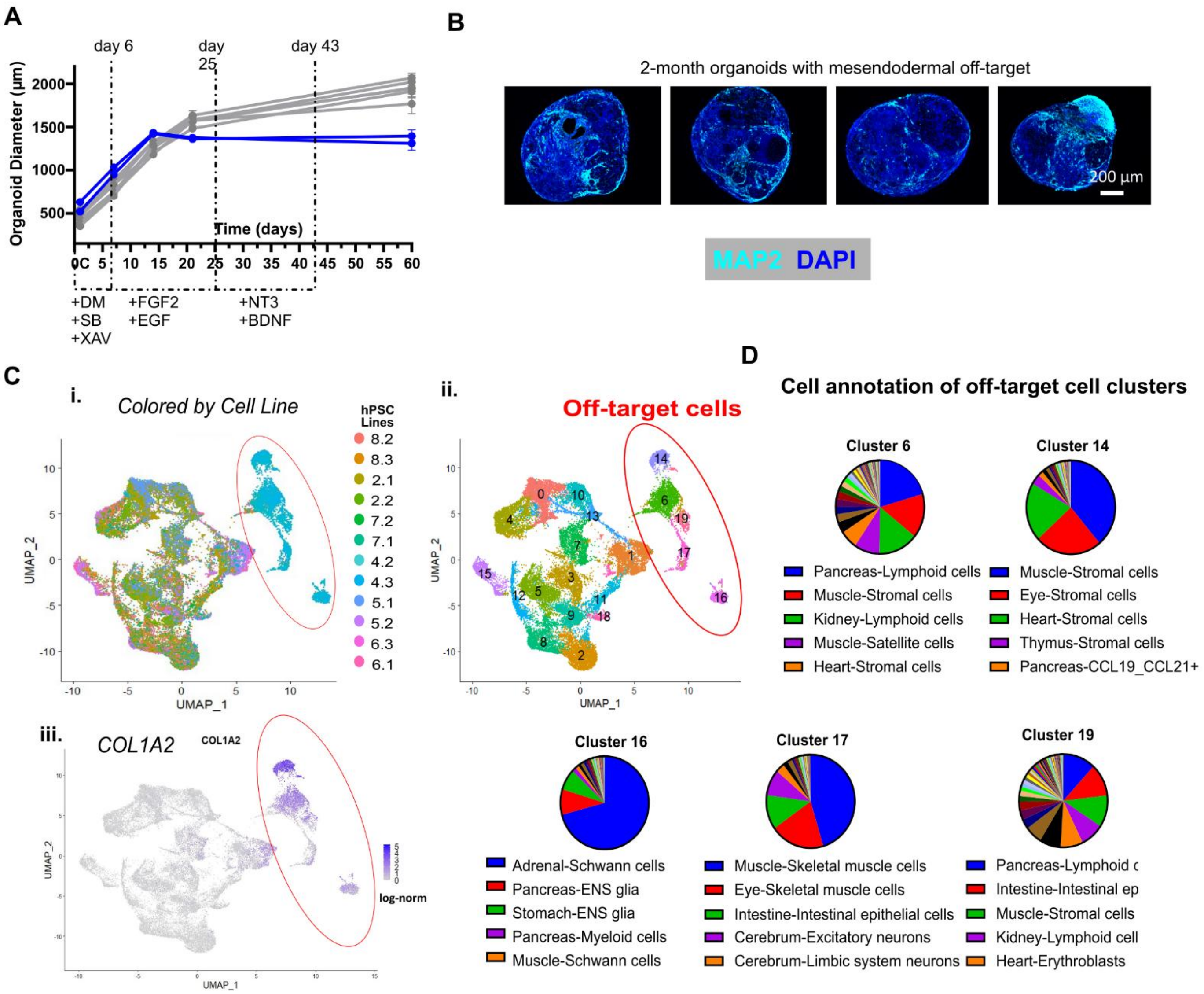


**Supplemental Figure S3. Impact of hPSC pluripotency on organoid production and mesendoderm contamination**. **S3A)** Mean organoid diameter over time in 96 slit-well plates from one experiment initiated from daily-fed mTeSR1 cultures (SD shown); blue lines denote donor #4 iPSC lines, which exhibited lower pluripotency markers prior to organoid production (5–12 organoids/line/timepoint; 12 lines). **S3B)** Representative immunofluorescence images of 2-month organoids from donor #4 lines (2 lines; 2–3 independent experiments). **S3C)** scRNA-seq of 2-month-old organoids generated from daily-fed hPSCs (12 lines; one experiment). Red ovals highlight off-target mesendodermal cell populations enriched in donor #4 organoids. These analyses identified cell populations lacking expression of neuronal genes (e.g., *MAPT*, *NEUROG2*) while expressing mesendoderm-associated genes (*COL1A1, COL1A2, TMEM119, DCN,* and *BGN*). (i) UMAP colored by line, (ii) Seurat clusters, (iii) *COL1A2* expression UMAP. **S3D)** Annotation against the human cell atlas of fetal gene expression[55] using the SingleR package[56] identified five cell clusters composed of various mesendodermal derivatives, including muscle, pancreas, kidney, lymphoid, adrenal, and Schwann cells. Cell-type composition of the five off-target clusters (clusters 6, 14, 16, 17, 19); labeled in S3Cii; ENS = enteric nervous system.

### Figure S4

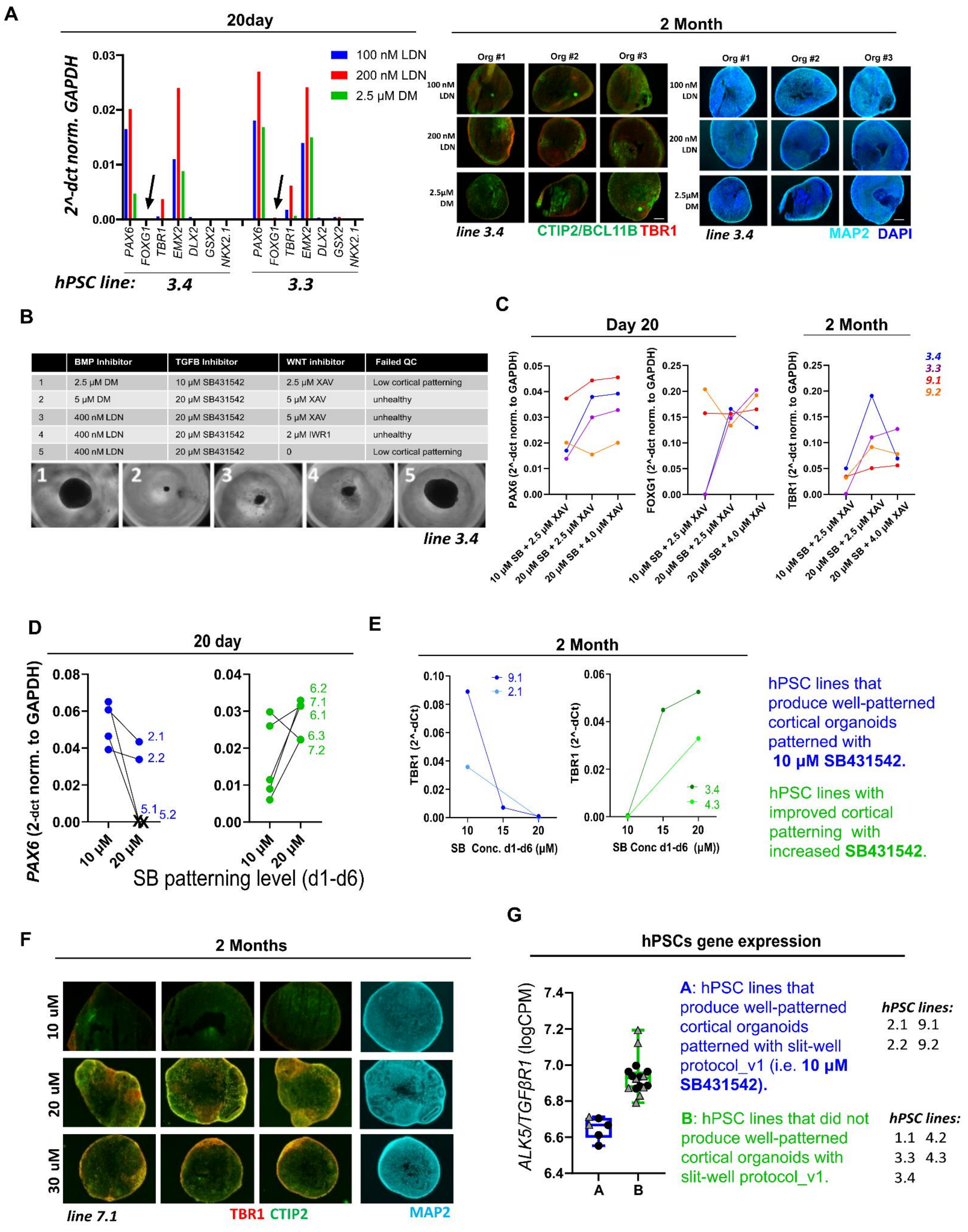


| | BMP Inhibitor | TGFB Inhibitor | WNT inhibitor | Failed QC |
|---|---|---|---|---|
| 1 | 2.5 µM DM | 10 µM SB431542 | 2.5 µM XAV | Low cortical patterning |
| 2 | 5 µM DM | 20 µM SB431542 | 5 µM XAV | unhealthy |
| 3 | 400 nM LDN | 20 µM SB431542 | 5 µM XAV | unhealthy |
| 4 | 400 nM LDN | 20 µM SB431542 | 2 µM IWR1 | unhealthy |
| 5 | 400 nM LDN | 20 µM SB431542 | 0 | Low cortical patterning |

**Supplemental Figure 4. Testing patterning molecules for cortical organoid production.**

**S4A)** Using 2 lines (3.3 and 3.4) that did not pattern well to cortex using slit-well protocol_v1, we tested several patterning molecule changes. Day 20 gene expression and representative images of 2-month organoid sections stained for deep-layer cortical markers TBR1 and CTIP2/BCL11B after BMP inhibition with varied dosing and/or inhibitors vs standard 2.5 µM dorsomorphin (DM); 3 pooled organoids/sample; 2 samples/condition; scale bar=200 µm. **S4B)** Patterning molecule combinations tested and outcomes with representative phase images of organoids in slit-wells. Removal of the WNT inhibitor XAV939 (2.5 µM; tankyrase inhibitor) impaired cortical patterning, consistent with prior work.[1] **S4C)** Day 20 and 2-month gene expression of organoids patterned with varying SB431542 (SB) and XAV939 (XAV) tested using lines that were successful (9.1, 9.2) or unsuccessful (3.3, 3.4) following slitwell protocol_v1 (shown in **Figure 1A**) (4 lines; 4 samples/condition; 3 pooled organoids/sample per datapoint). **S4D)** *PAX6* expression in day 20 organoids patterned with 10 vs 20 µM SB431542 (9 lines, 6 donors; 3 pooled organoids/datapoint); 'X' denotes culture lost due to low viability (lines 5.1, 5.2), while lower *PAX6* levels in other lines (2.1, 2.2) indicate suboptimal patterning. **S4E)** *TBR1* expression in 2-month organoids patterned with SB431542 (d1–d6): lines optimized at 10 µM (blue) vs 20 µM (green) (4 lines; 4 samples/condition; 3 pooled organoids/datapoint; 2 independent experiments). **S4F)** Representative 2-month sections stained for MAP2, TBR1, and CTIP2/BCL11B after patterning with 10, 20, or 30 µM SB431542 (d1–d6); 3 organoids/condition; scale bar=200 µm. **S4G)** *ALK5* (TGFβ receptor) expression is lower in hPSCs that pattern well with 10 µM SB431542 (blue; 4 lines, 2 repeated; 6 samples) vs lines that pattern poorly (green; 6 lines, each repeated 2–3 times; 16 samples); 3 independent experiments; CPM, counts per million; culture method indicated by shape (triangle, daily fed; circle, FGF2DISC).

### Figure S5

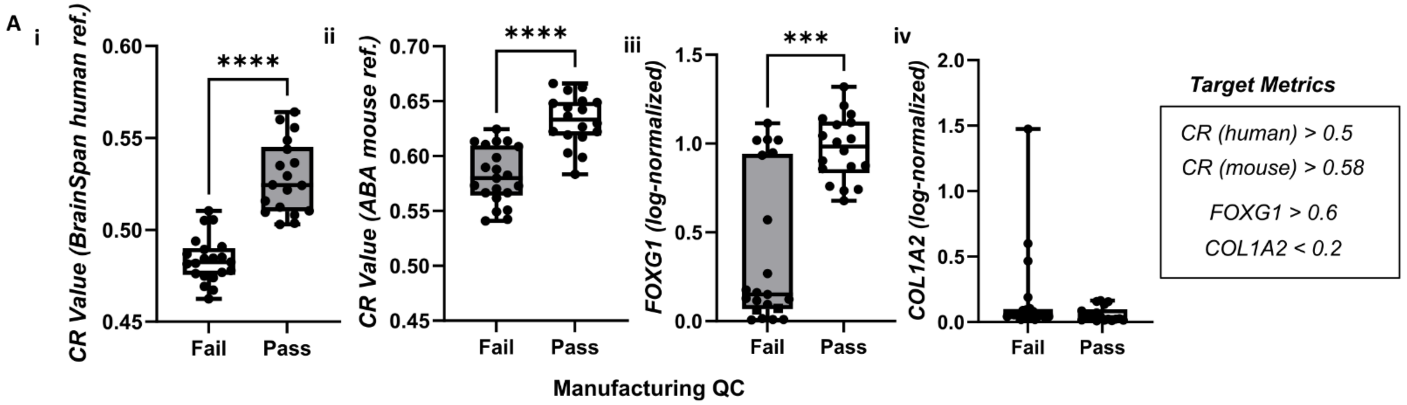


B

$$\text{CortiCOSI} = 0.5\left(\frac{x-\mu_x}{\sigma_x}\right) + 0.25\left(\frac{y-\mu_y}{\sigma_y}\right) - 0.25\left(\frac{z-\mu_z}{\sigma_z}\right)$$

**x = CR**: all cells (calculated using VoxHunt and a reference dataset)
**y = *FOXG1* level**: all cells (log-Norm)
**z = *COL1A2* level**: all cells (log-Norm)

**Figure S5. scRNA-seq–based quality assessment of cortical organoids and CortiCOSI metric development.** **S5A)** Cortical organoid quality assessed by scRNA-seq metrics. i,ii) the CR identity values measured by VoxHunt. (i) ABA mouse reference[1], unpaired two-tailed t-test, ****p <0.0001; ii) Human BrainSpan reference[2], unpaired two-tailed t-test, ****p <0.0001; iii) *FOXG1* expression (log-normalized); Mann-Whitney U test (two-tailed), ***p=0.0001; (iv) *COL1A2* expression (log-normalized). **S5B**) Equation for CortiCOSI that integrates all three metrics with weighted z-scores; μ=mean; σ=standard deviation.

### Figure S6

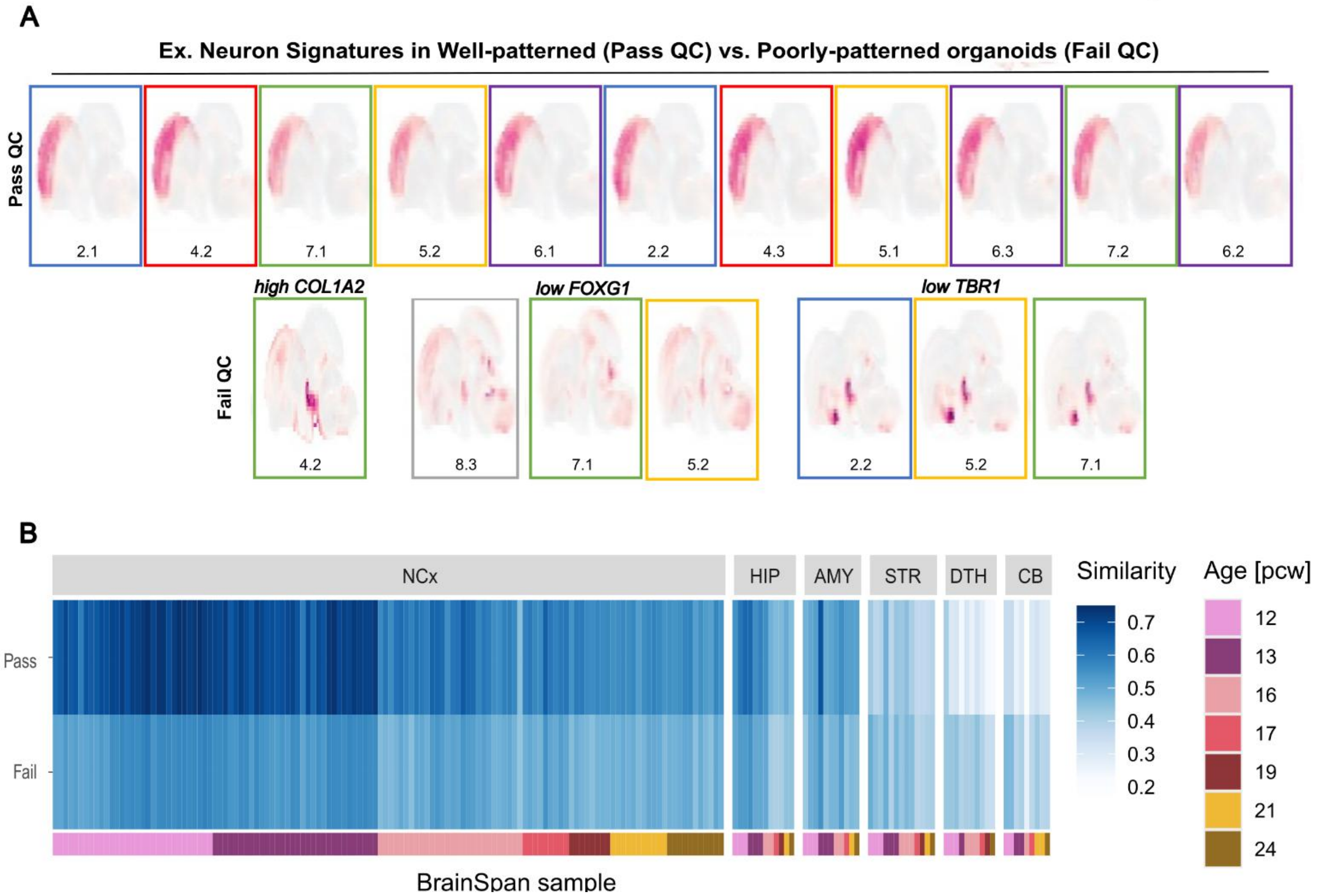


**Supplemental Figure 6. Neuronal cortical identity projections.**

**S6. S6A,B)** Excitatory neuron signature projections onto developing brain reference datasets, stratified by pass or fail manufacturing QC. **S6A)** VoxHunt projections from individual organoid samples (21 representative images of 39 total samples). **S6B)** Heatmap showing correlation with the BrainSpan human reference.

#### Figure S7

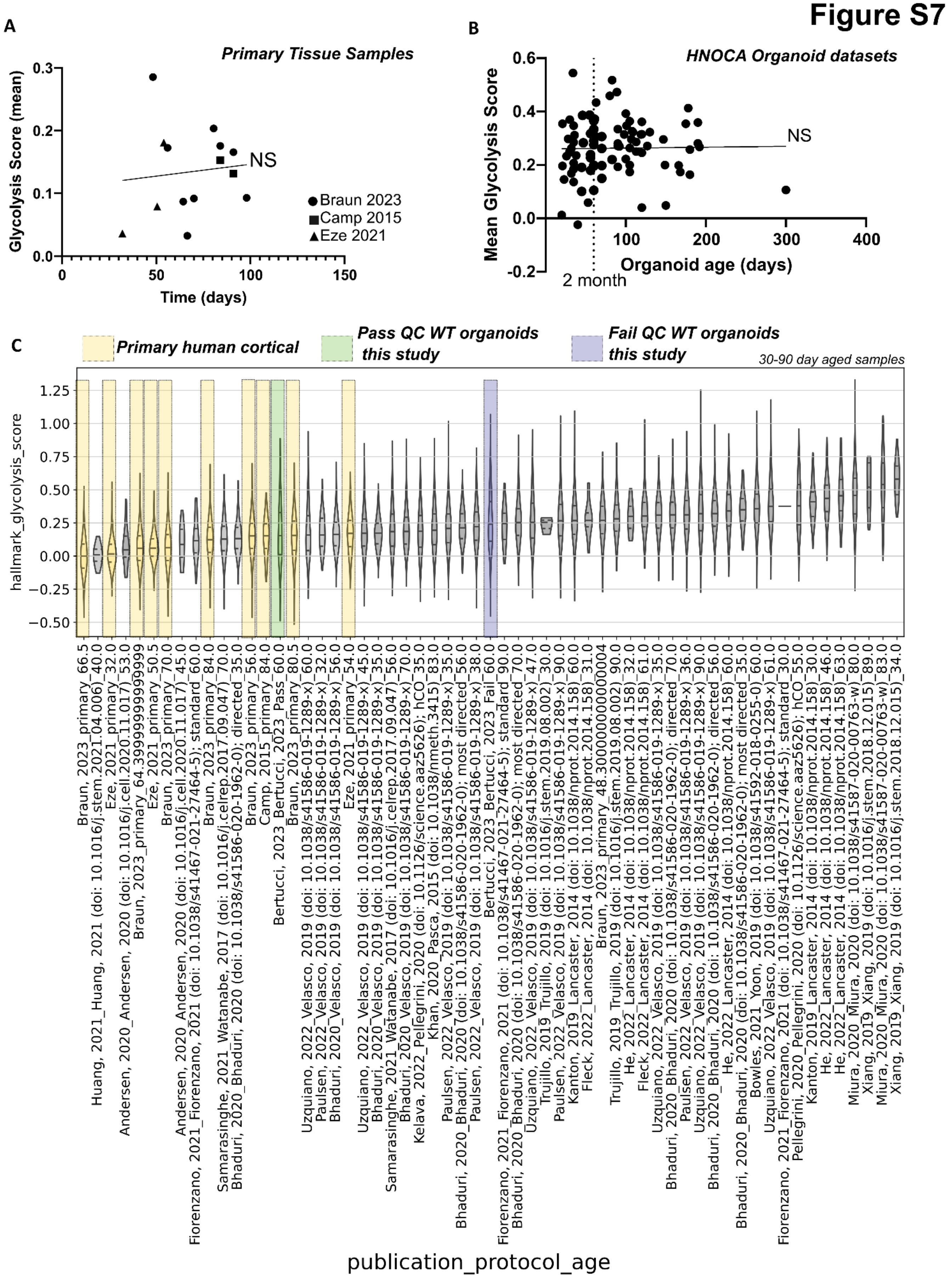

A
Primary Tissue Samples
Glycolysis Score (mean)
0.0
0.1
0.2
0.3
NS
Braun 2023
Camp 2015
Eze 2021
0
50
100
150
Time (days)
B
HNOCA Organoid datasets
Mean Glycolysis Score
-0.2
0.0
0.2
0.4
0.6
NS
100
200
300
400
Organoid age (days)
2 month
C
Primary human cortical
Pass QC WT organoids this study
Fail QC WT organoids this study
30-90 day aged samples
hallmark_glycolysis_score
−0.50
−0.25
0.00
0.25
0.50
0.75
1.00
1.25
Braun, 2023_primary_66.5
Huang, 2021_Huang, 2021 (doi: 10.1016/j.stem.2021.04.006)_40.0
Eze, 2021_primary_32.0
Andersen, 2020_Andersen, 2020 (doi: 10.1016/j.cell.2020.11.017)_53.0
Braun, 2023_primary_64.39999999999999
Eze, 2021_primary_50.5
Braun, 2023_primary_70.0
Andersen, 2020_Andersen, 2020 (doi: 10.1016/j.cell.2020.11.017)_45.0
Fiorenzano, 2021_Fiorenzano, 2021 (doi: 10.1038/s41467-021-27464-5); standard_60.0
Braun, 2023_primary_84.0
Samarasinghe, 2021_Watanabe, 2017 (doi: 10.1016/j.celrep.2017.09.047)_70.0
Bhaduri, 2020_Bhaduri, 2020 (doi: 10.1038/s41586-020-1962-0); directed_35.0
Braun, 2023_primary_56.0
Camp, 2015_primary_84.0
Bertucci, 2023_Bertucci, 2023_Pass_60.0
Braun, 2023_primary_80.5
Uzquiano, 2022_Velasco, 2019 (doi: 10.1038/s41586-019-1289-x)_60.0
Paulsen, 2022_Velasco, 2019 (doi: 10.1038/s41586-019-1289-x)_32.0
Bhaduri, 2020_Velasco, 2019 (doi: 10.1038/s41586-019-1289-x)_56.0
Eze, 2021_primary_54.0
Uzquiano, 2022_Velasco, 2019 (doi: 10.1038/s41586-019-1289-x)_45.0
Bhaduri, 2020_Velasco, 2019 (doi: 10.1038/s41586-019-1289-x)_35.0
Samarasinghe, 2021_Watanabe, 2017 (doi: 10.1016/j.celrep.2017.09.047)_56.0
Bhaduri, 2020_Velasco, 2019 (doi: 10.1038/s41586-019-1289-x)_70.0
Kelava, 2022_Pellegrini, 2020 (doi: 10.1126/science.aaz5626); hCO_35.0
Khan, 2020_Pasca, 2015 (doi: 10.1038/nmeth.3415)_83.0
Paulsen, 2022_Velasco, 2019 (doi: 10.1038/s41586-019-1289-x)_35.0
Bhaduri, 2020_Bhaduri, 2020 (doi: 10.1038/s41586-020-1962-0); most directed_56.0
Paulsen, 2022_Velasco, 2019 (doi: 10.1038/s41586-019-1289-x)_38.0
Bertucci, 2023_Bertucci, 2023_Fail_60.0
Fiorenzano, 2021_Fiorenzano, 2021 (doi: 10.1038/s41467-021-27464-5); standard_90.0
Bhaduri, 2020_Bhaduri, 2020 (doi: 10.1038/s41586-020-1962-0); most directed_70.0
Uzquiano, 2022_Velasco, 2019 (doi: 10.1038/s41586-019-1289-x)_47.0
Trujillo, 2019_Trujillo, 2019 (doi: 10.1016/j.stem.2019.08.002)_30.0
Paulsen, 2022_Velasco, 2019 (doi: 10.1038/s41586-019-1289-x)_90.0
Kanton, 2019_Lancaster, 2014 (doi: 10.1038/nprot.2014.158)_60.0
Fleck, 2022_Lancaster, 2014 (doi: 10.1038/nprot.2014.158)_31.0
Braun, 2023_primary_48.300000000000004
Trujillo, 2019_Trujillo, 2019 (doi: 10.1016/j.stem.2019.08.002)_90.0
He, 2022_Lancaster, 2014 (doi: 10.1038/nprot.2014.158)_32.0
Fleck, 2022_Lancaster, 2014 (doi: 10.1038/nprot.2014.158)_61.0
Uzquiano, 2022_Velasco, 2019 (doi: 10.1038/s41586-019-1289-x)_35.0
Bhaduri, 2020_Bhaduri, 2020 (doi: 10.1038/s41586-020-1962-0); directed_70.0
Paulsen, 2022_Velasco, 2019 (doi: 10.1038/s41586-019-1289-x)_36.0
Uzquiano, 2022_Velasco, 2019 (doi: 10.1038/s41586-019-1289-x)_90.0
Bhaduri, 2020_Bhaduri, 2020 (doi: 10.1038/s41586-020-1962-0); directed_56.0
He, 2022_Lancaster, 2014 (doi: 10.1038/nprot.2014.158)_60.0
Bhaduri, 2020_Bhaduri, 2020 (doi: 10.1038/s41586-020-1962-0); most directed_35.0
Bowles, 2021_Yoon, 2019 (doi: 10.1038/s41592-018-0255-0)_60.0
Uzquiano, 2022_Velasco, 2019 (doi: 10.1038/s41586-019-1289-x)_61.0
Fiorenzano, 2021_Fiorenzano, 2021 (doi: 10.1038/s41467-021-27464-5); standard_30.0
Pellegrini, 2020_Pellegrini, 2020 (doi: 10.1126/science.aaz5626); hCO_55.0
Kanton, 2019_Lancaster, 2014 (doi: 10.1038/nprot.2014.158)_30.0
He, 2022_Lancaster, 2014 (doi: 10.1038/nprot.2014.158)_46.0
He, 2022_Lancaster, 2014 (doi: 10.1038/nprot.2014.158)_63.0
Miura, 2020_Miura, 2020 (doi: 10.1038/s41587-020-00763-w)_80.0
Xiang, 2019_Xiang, 2019 (doi: 10.1016/j.stem.2018.12.015)_89.0
Miura, 2020_Miura, 2020 (doi: 10.1038/s41587-020-00763-w)_83.0
Xiang, 2019_Xiang, 2019 (doi: 10.1016/j.stem.2018.12.015)_34.0
publication_protocol_age

**Supplemental Figure S7. Metabolic stress profiling across neurons in cortical organoids and human tissue. S7A,B)** Glycolysis stress score plotted across developmental time in human cortical tissue datasets[3-5] and HNOCA organoid datasets[6] (right); the stress score does not correlate with human developmental timepoint or organoid age; linear regression, NS=not significant. **S7C)** Neuron stress scores across the HNOCA organoid dataset[6] (30–90 days), human cortical tissue[2-4], and datasets generated in this study, stratified by QC pass/fail status are highlighted. Lines within violin plots indicate quartiles.

### Figure S8

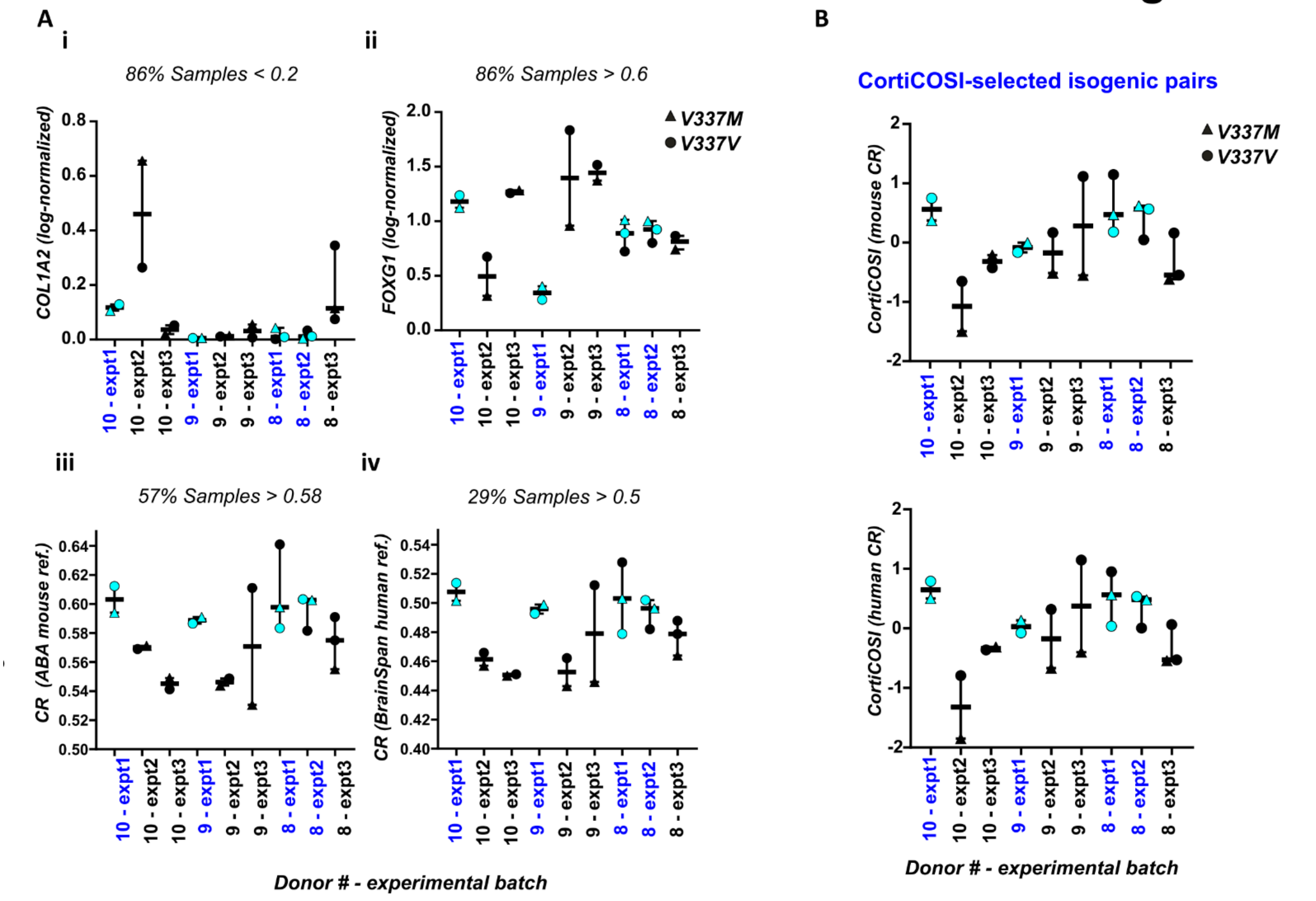


**Figure S8. CortiCOSI-based assessment of quality and variability in V337M organoids.**

**S8A)** scRNA-seq assessment of cortical organoid quality in *MAPT* V337M organoids generated in dishes[7] assessed by isogenic set; 3 isogenic sets (2-3 lines per set), 3 experiments. i) *COL1A2* expression, ii) *FOXG1* expression ii) CR value (mouse reference) iv) CR value (human reference). **S7B**) CortiCOSI scores to assess inter-sample variation across isogenic pairs. Blue notation indicates isogenic sets selected to be the highest quality and most consistent.

**Figure S9**

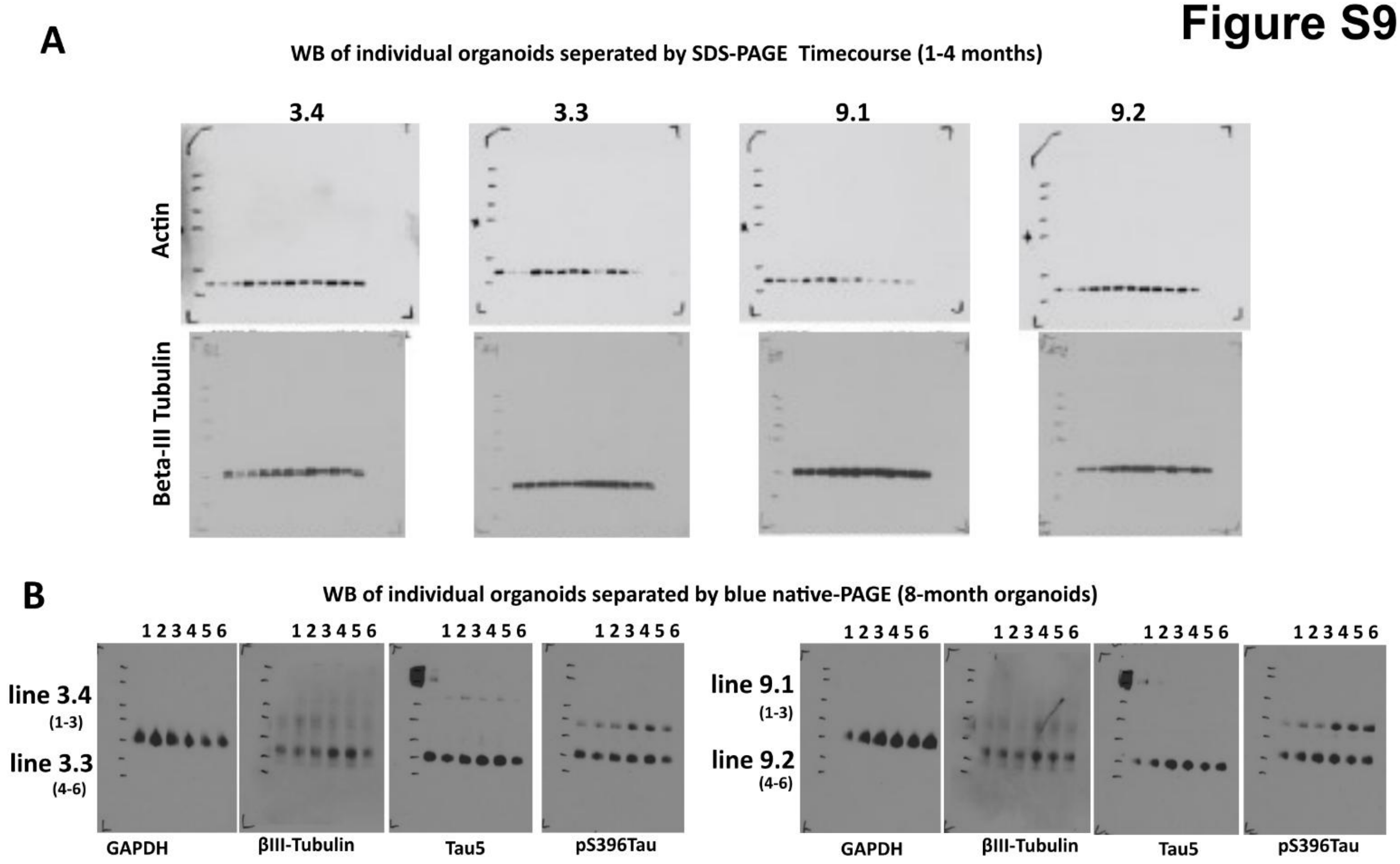


**Supplemental Figure 9. Raw and uncropped western blot images.**

**S9A)** Lysates from individual organoids collected at 1, 2, 3, and 4 months were analyzed by WB, separated by denaturing SDS-PAGE; see **Figure S1C** legend for sample numbers. **S9B)** Lysates from individual organoids collected at 8 months (250 days) were analyzed by WB, separated by non-denaturing BN-PAGE; see **Figure 6D** legend for sample numbers.

1 Fleck, J. S. *et al.* Resolving organoid brain region identities by mapping single-cell genomic data to reference atlases. *Cell Stem Cell* **28**, 1148-1159 e1148 (2021). https://doi.org/10.1016/j.stem.2021.02.015
2 Kang, H. J. *et al.* Spatio-temporal transcriptome of the human brain. *Nature* **478**, 483-489 (2011). https://doi.org/10.1038/nature10523
3 Braun, E. *et al.* Comprehensive cell atlas of the first-trimester developing human brain. *Science* **382**, eadf1226 (2023). https://doi.org/10.1126/science.adf1226
4 Camp, J. G. *et al.* Human cerebral organoids recapitulate gene expression programs of fetal neocortex development. *Proc Natl Acad Sci U S A* **112**, 15672-15677 (2015). https://doi.org/10.1073/pnas.1520760112
5 Eze, U. C., Bhaduri, A., Haeussler, M., Nowakowski, T. J. & Kriegstein, A. R. Single-cell atlas of early human brain development highlights heterogeneity of human neuroepithelial cells and early radial glia. *Nat Neurosci* **24**, 584-594 (2021). https://doi.org/10.1038/s41593-020-00794-1
6 He, Z. *et al.* An integrated transcriptomic cell atlas of human neural organoids. *Nature* **635**, 690-698 (2024). https://doi.org/10.1038/s41586-024-08172-8
7 Bowles, K. R. *et al.* ELAVL4, splicing, and glutamatergic dysfunction precede neuron loss in MAPT mutation cerebral organoids. *Cell* **184**, 4547-4563 e4517 (2021). https://doi.org/10.1016/j.cell.2021.07.003
